# Confidence over competence: Real-time integration of social information in human continuous perceptual decision-making

**DOI:** 10.1101/2024.08.19.608609

**Authors:** Felix Schneider, Antonino Calapai, Roger Mundry, Raymundo Báez-Mendoza, Alexander Gail, Igor Kagan, Stefan Treue

## Abstract

Human perceptual decision-making is susceptible to social influences. To determine if and how individuals opportunistically integrate real-time social information about noisy stimuli into their judgment, we tracked perceptual accuracy and confidence in social (dyadic) and non-social (solo) settings using a novel continuous perceptual report (CPR) task with peri-decision wagering. In the dyadic setting, most participants showed a higher degree of perceptual confidence. In contrast, average accuracy did not improve compared to solo performance. Underlying these net effects, partners in a dyad exhibit mutual convergence of accuracy and confidence, benefiting less competent or confident individuals, at the expense of the better-performing partner. In conclusion, real-time social information asymmetrically shapes human perceptual decision-making, with most dyads expressing more confidence without a matching gain in overall competence.

## Introduction

To navigate dynamic and uncertain environments, people continuously gather and evaluate sensory evidence. This ongoing perceptual decision process is shaped not only by competence – the accuracy of percept – but also confidence: the belief in that accuracy, formed through metacognitive evaluation of current and past decisions. Confidence plays a pivotal role because it governs how individuals act on their perceptions, whether committing to a choice, revising it, or seeking more information (Fleming and Lau, 2014; Kepecs and Mainen, 2012; Yeung and Summerfield, 2012).

In social situations, people routinely adjust their decisions and confidence in response to others’ behavior, feedback, or consensus – even when their own sensory input remains unchanged (Bahrami et al., 2010; Bang et al., 2017; Esmaily et al., 2023; Pescetelli et al., 2016; Pescetelli and Yeung, 2022). These adjustments reflect susceptibility to both informational and normative social influences (Frith and Singer, 2008; Takagaki and Krug, 2020; Terenzi et al., 2021; Toelch and Dolan, 2015; Van Den Bos et al., 2013). For instance, by integrating the perceptual report of a partner with one’s own subjective experience, the quality of perceptual judgments can be optimized and result in a collective benefit, but only under certain conditions (Bahrami et al., 2010, 2012a; Baumgart et al., 2020). Social information can speed up decision-making by reducing exploration time, but it can also introduce biases and false beliefs (Bang and Frith, 2017). For example, social conformity biases information uptake towards majority choices (Germar et al., 2016; Toelch et al., 2018).

Expressions of confidence in particular have been shown to exert a major influence during social exchange. Judging competence of others is difficult because it requires performance monitoring over time, and accuracy and confidence measures generally covary (Baumgart et al., 2020; Khalvati et al., 2021; Pescetelli et al., 2016). Therefore, people often use confidence signals of a partner as a proxy for assessing competence of others (Bang and Frith, 2017). Furthermore, confidence signals could serve as a channel for social information transfer. For instance, the expressed confidence range can adapt to specific social partners to achieve more optimal group decisions (Bang et al., 2017). Access to choices coupled with the confidence reports of another player has been shown to asymmetrically modulate individual decisions: compared to solo performance, dyadic agreement caused greater improvement in accuracy and confidence levels than disagreement reduced these measures (Pescetelli et al., 2016).

While these and other studies have illuminated the interplay of individual decisions and social influences, they relied on paradigms that treat perceptual decisions and confidence judgments as discrete and sequential – limiting our understanding of how these highly intertwined processes unfold during continuous, real-time decision-making. First, stimuli are typically presented as serial, isolated events rather than ongoing, temporally correlated streams, which are more representative of real-world environments. Second, many studies employed trial-based, discrete paradigms in which perceptual decisions and confidence reports are temporally separated. This separation introduces the possibility that metacognitive judgments draw on information not available at the time of the perceptual experience itself (Navajas et al., 2016; Yeung and Summerfield, 2012). Third, decisions were constrained to binary alternatives, e.g., left vs right (Esmaily et al., 2023; Kiani and Shadlen, 2009) or first vs second interval (Bahrami et al., 2010; Pescetelli and Yeung, 2022), precluding a more graded expression of perceptual experience. Fourth, most studies imposed rigid structure with individual choices preceding social exchange and joint decision-making, assessing interdependent dyadic performance (Bahrami et al., 2010; Bang et al., 2017; Baumgart et al., 2020). To understand how accuracy and confidence evolve during the course of the dynamic decision-making and how both factors are shaped by the availability of social information, novel experimental methods allowing access to continuous reports are needed.

Recent work began to demonstrate the influence of continuous information exchange during dyadic co-action, although still in a two-alternative and serial design. Dynamic, mutually visible confidence reports elicited greater increase of perceptual confidence during agreement and less reduction during disagreement, compared to static reports (Pescetelli and Yeung, 2020), and also resulted in a confidence alignment between partners (Pescetelli and Yeung, 2022). Emergent continuous psychophysics approaches further capture dynamics of perception, quantifying sensorimotor and cognitive processes via continuous perceptual report (CPR) tasks (Bonnen et al., 2017, 2015; Huk et al., 2018; Straub and Rothkopf, 2022). As CPRs can be displayed continuously, they are particularly useful to elucidate interactions that unfold over time, such as the integration of noisy sensory and social information.

Motivated by this goal, we investigated how real-time, unconstrained social information influences the continuous expression of perceptual accuracy and confidence during decision-making about ambiguous visual stimuli. We developed a continuous “peri-decision” wagering paradigm, allowing participants to signal their percept and associated confidence in real-time using simple visual cues observable by others. This contrasts with previous approaches that relied on post-decision confidence measures – such as numerical ratings (Boldt and Yeung, 2015; Fleming et al., 2010), post-decision wagering (Moreira et al., 2018; Persaud et al., 2007), or opt-out/decline response options (Gail et al., 2004; Hanks and Summerfield, 2017; Khalvati et al., 2021; Kiani and Shadlen, 2009; Komura et al., 2013; Smith, 1997). Notably, our “co-action” design did not enforce payoff interdependence between players: they were free to monitor and opportunistically incorporate their partner’s report, without being compelled to do so.

This approach allowed us to address three key questions: (1) Do humans express perceptual confidence in real-time? (2) Does social context influence both perceptual accuracy and confidence expression? (3) How does the competence and the confidence of a social partner shape the integration of perceptual and social evidence? We hypothesized that participants continuously adjust confidence based on moment-to-moment sensory evaluation and selectively integrate their partner’s cues to interpret ambiguous stimuli, resulting in more accurate and confident reports. We further predicted that social benefit will depend on relative competence, with lower-performing individuals gaining more from interaction with a more skilled partner.

## Results

To assess the integration of social information, we developed a dynamic perceptual decision paradigm (**Figure 1A,B**, **Supplementary Figure 1A** and **Supplementary Video 1**, Continuous Perceptual Report task, ‘CPR’, see Methods and Glossary) that enables continuous “peri-decision” wagering on the accuracy of perceptual judgments about the direction of a noisy random dot pattern (RDP). The RDP direction and coherence changed frequently, resulting in successive “stimulus states” (**Figure 1C**). Participants used a joystick to signal their perceived direction (angle) and confidence (joystick tilt, the deviation from the central position). Their task was to maximize the monetary reward score by following the direction of the RDP as accurately as possible, so that their cursor (partial circle, ‘response arc’) would overlap with occasionally presented small reward targets appearing in the direction of the coherent motion (**Supplementary Figure 1B**). Tilting the joystick away from the center shortened the arc length, making it harder to hit the target and obtain the reward. But at the same time, larger tilt could result in a larger reward, because the reward score was calculated as the product of report accuracy and tilt (but was zero if the response arc did not hit the target, **Figure 1D**). This reward scheme allowed us to elicit continuous peri-decision wagering, by awarding large rewards for accurate and confident reports, small rewards for less accurate and/or less confident reports, and omitting rewards for inaccurate and overconfident reports.

**Figure 1:**
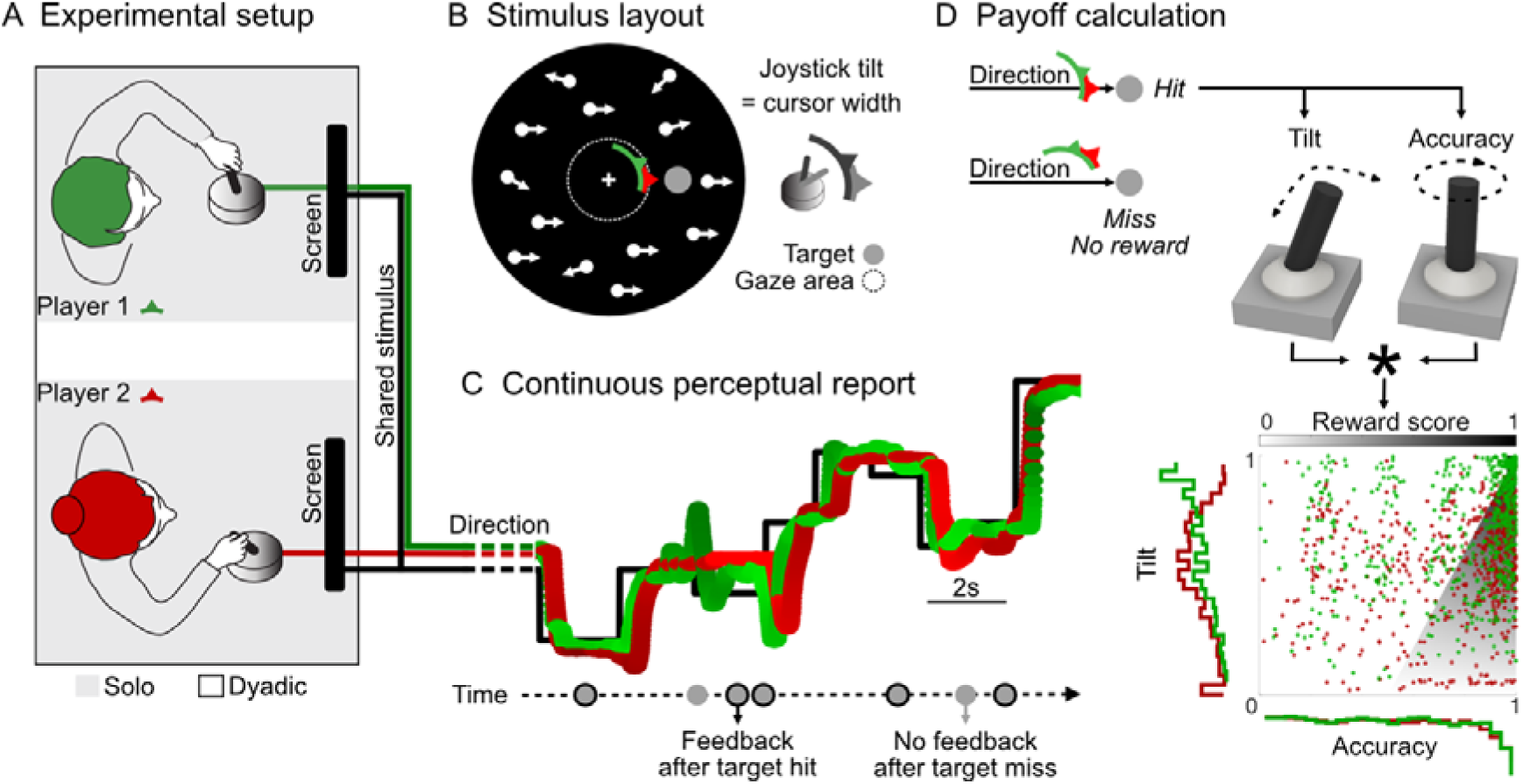
Continuous perceptual report task. **(A)** Experimental setup: Two participants sat in adjacent experimental booths. Subjects played a motion tracking game with a joystick either alone (‘solo’) or together with a partner (‘dyadic’, mixed order: see **Supplementary Figure 1A**). In dyadic experiments, subjects watched the shared visual stimulus on a screen. Joystick responses of both players as well as visual feedback were mutually visible in real-time. **(B)** Random dot pattern (‘RDP’) with circular aperture and blacked out central fixation area was continuously presented for intervals of about 1 minute. Subjects were instructed to look at the central fixation cross. Joystick-controlled cursors (‘response arcs’, color-coded) are located at the edge of the fixation area. The stimulus motion signal was predictive of the location of behaviorally relevant reward targets (gray disc). Alignment of the cursor arc with the target resulted in target collection (‘hit’) and reward. The joystick tilt was linked to size of the cursor: less tilt – wide; more tilt - narrow. **(C)** Example trace of stimulus motion signal and dyadic responses: the stimulus was frequently changing in motion direction and coherence level. Participants tracked the stimulus direction with the joystick to obtain rewards. Darker hues indicate less joystick tilt. Target presentation occurred in pseudorandom time intervals (see **Supplementary Figure 1B**). The visual and auditory feedback representing the reward score was provided after each target hit. **(D)** Top: Upon target hit, the reward score was based on joystick tilt and accuracy, individually for each player. Bottom: Each dot corresponds to the accuracy-tilt combination during target presentation. The shaded greyscale background illustrates the non-zero reward map. The non-shaded area denotes missed targets with no reward (reward score = 0). The accuracy and tilt responses of both dyadic players are summarized with the histograms (color-coded). Positive trend of reward scores, shown in **Supplementary Figure 1C**, indicates perceptual learning over time.

Participants played the game in different experimental settings: alone (*solo)* or simultaneously with a partner (*dyadic*). In dyadic experiments, the joystick direction and tilt of both players are continuously presented to both participants. Here, we define social information as the perceptual report of the partner.

### Humans express perceptual confidence in real-time through peri-decision wagering

Confidence measures have been shown to scale with evidence accumulation time and perceptual performance (Balsdon et al., 2021, 2020; Kiani and Shadlen, 2009). Our task design intends to capture perceptual confidence via real-time wagering behavior. In this section, we verify that participants used peri-decision wagering while playing the CPR game.

More accurate participants showed more joystick tilt (Pearson’s correlation of average accuracy and tilt, n = 38 subjects, r = 0.62, p<0.0001). Participants increased their hit rate (fraction of successfully acquired targets) and joystick tilt, while reducing their angular error (i.e., improving accuracy), with higher motion coherence (**Figure 2A**). Their response lag after an RDP direction change was on average 643 ms ± 79 ms (Mean ± IQR, coherence pooled; **Figure 2B-C**, **Supplementary Figure 1D**), with higher motion coherence causing shorter stimulus following responses. Low RDP coherence resulted in a breakdown of motion tracking, increasing the variance of cross-correlation peaks (**Supplementary Figure 1D**). Participants also increased their tracking accuracy and joystick tilt within individual stimulus states, except at very low coherences (**Figure 2D**, Linear regression of the average time course in each participant, Accuracy: Mean slope = 4.8835e-04; Wilcoxon signed rank test against zero: n = 38 subjects, Z = 5.3731, p < 0.0001; Tilt: Mean slope = 1.0366e-04; Wilcoxon signed rank test against zero: n = 38 subjects, Z = 3.2993, p < 0.001), indicating a continuous evidence accumulation during stable stimulus epochs. Changes in accuracy preceded changes in joystick tilt (**Figure 2E**, Cross-correlation, Mean lag = 339.9 ms, Wilcoxon signed rank test against zero: n = 38 subjects, Z = 5.3733, p < 0.0001). These patterns reflect the dynamics of evidence integration, with faster and more reliable responses driving higher levels of confidence.

**Figure 2:**
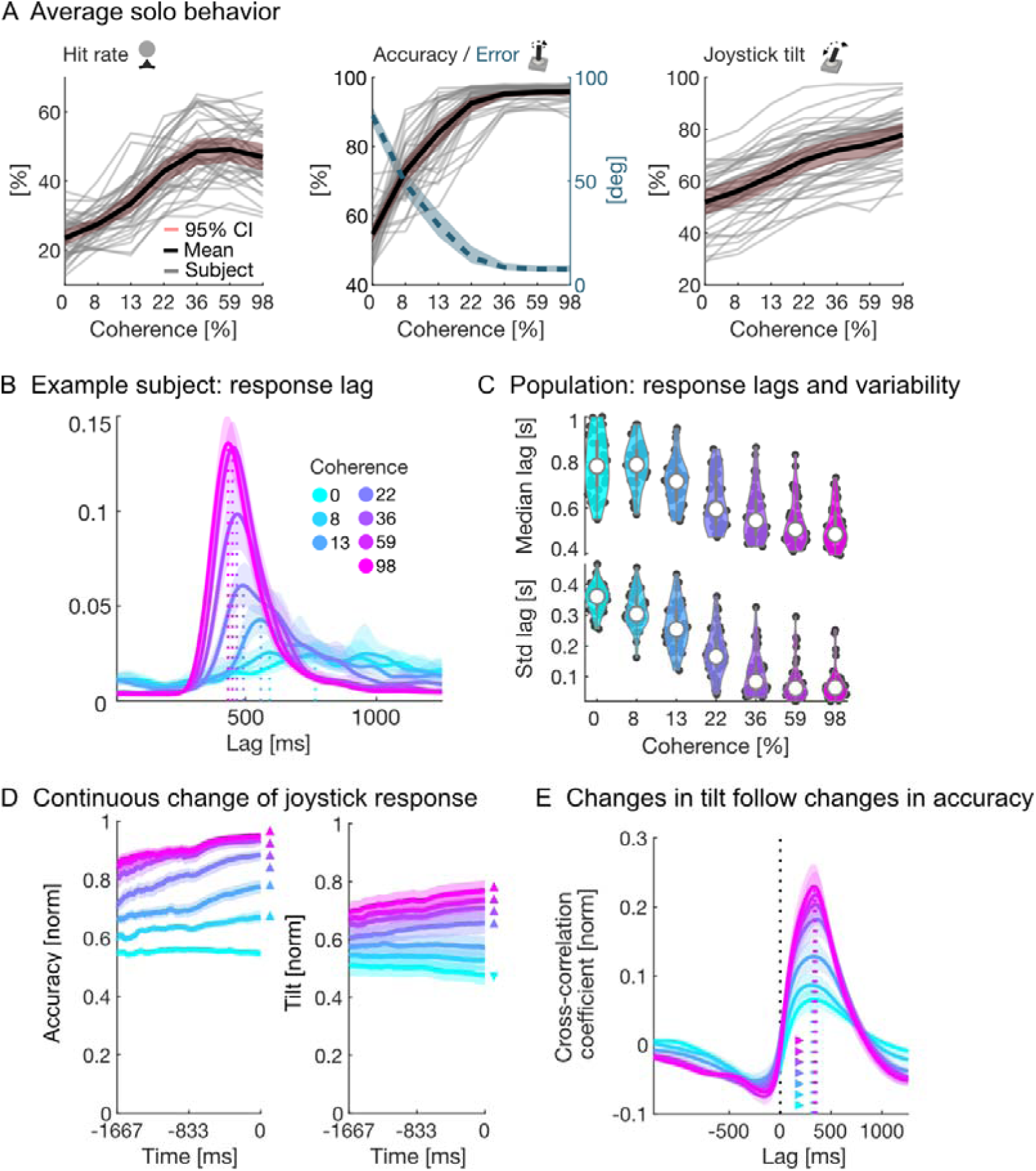
Solo behavior during the continuous perceptual report. **(A)** Coherence-dependent modulation of hit rates (left), accuracy (center, shown with absolute, target-aligned angular joystick error as dashed blue line) and state-aligned radial joystick tilt (right). Gray lines show averages for individual participants. Red shading illustrates the 99% confidence intervals of the mean across participants. Bold, black lines show the mean across the population. Data were first averaged within-subject, before pooling coherence conditions across-subjects. **(B)** Estimation of the response lag after a stimulus direction change with a cross-correlation between stimulus and joystick signal for an example subject (dashed lines). Low coherence levels resulted in a breakdown of response reliability, indicated by the low cross-correlation coefficients (see also **Supplementary Figure 1D**). Here and in other panels, stimulus coherence is color-coded. **(C)** Top: average population response lag. Black data points show individual data; white dot displays population median. Bottom: variability of the population lags, displayed by the standard deviation. **(D)** Average time course of joystick accuracy (left) and tilt (right) during a stable stimulus period aligned to the next stimulus direction change at time point 0, across all subjects. Here, the first 500 ms of each stimulus state as well as all samples after the first target appearance were excluded before averaging. The shaded background illustrates the 95% confidence interval of the mean. We calculated the slopes of the average time course and tested if they were significantly different from zero across subjects (n = 38, Bonferroni-corrected). Statistics is illustrated with color-coded triangles, indicating which coherence condition has a significant slope and in which direction (triangles point up for positive slopes). **(E)** Cross-correlation between the mean-detrended time course of joystick tilt and accuracy, indicating that changes in tilt follow changes in accuracy within half a second. Peak cross-correlation coefficients are marked with a dashed line. Statistics is illustrated with the color-coded triangles, indicating which coherence condition has a significantly shifted from 0 cross-correlation peak across subjects (n = 38, Bonferroni-corrected).

Despite high inter-subject variability, we found that motion coherence had a robust impact on all behavioral response measures: joystick accuracy, tilt and hit rates (Linear mixed effects models – see **Supplementary Table 1** to **Supplementary Table 4**). This suggests that participants adapted their responses to varying stimulus difficulty. We further demonstrate that hit rates, while generally increasing with motion coherence, dropped for 24 of 38 participants (63%) during the most salient RDP coherence, suggesting overconfident or risk-seeking joystick placement.

To directly assess if subjects used joystick tilt as a proxy for perceptual confidence, we adapted the metacognitive performance analysis of the area under the receiver operating characteristics curve (AUC) for confidence ratings to continuous joystick responses (Fleming and Lau, 2014; Maniscalco and Lau, 2014, 2012). We used the distributions of joystick response measurements to infer whether there is a relationship between response accuracy and tilt, separately for each RDP coherence level. To that end, we median-split the joystick responses into high and low tilt distributions. We then analyzed the AUC to quantify whether the accuracy of these distributions was different. High accuracy was, indeed, related to more joystick tilt, suggesting an ongoing metacognitive assessment of the perceptual report that is reflected in the tilt of the joystick (**Supplementary Figure 2**). Thus, participants optimized joystick placement in both accuracy and tilt, which provides evidence for real-time peri-decision wagering and links joystick tilt to perceptual confidence. Importantly, we ruled out that the short-term fluctuations of instantaneous coherence or random differences in average coherence between stimulus states could explain the variations in joystick tilt within each nominal coherence level. Furthermore, the gradual tilt increase within stable stimulus epochs indicates that this response dimension was dissociated from the fluctuations of motion coherence (**Figure 2D**).

In summary, solo CPRs indicate that participants actively wager on their own percept. As joystick tilt was the only response dimension that could be chosen freely via metacognitive assessment of the current and past decisions, we treat it as a proxy measure of subjective perceptual confidence. Usage and range of this response parameter varied widely between participants, suggesting individual confidence ranges. These findings indicate that our CPR game makes it possible to continuously assess participants’ perceptual processes and associated confidence. Next, we examine whether and to what extent participants incorporated the perceptual report of a second player into their own decision-making.

### Social setting changes perceptual accuracy and confidence during real-time decision-making

Previous studies have shown that, under certain conditions, two participants are more successful in perceptual decision-making than the more competent player on its own (Bahrami et al., 2012b; Pescetelli and Yeung, 2022). We asked whether participants performed the CPR task better when a second player was playing along, and if so, whether differences between participants in perceptual competence or confidence were the driving forces behind any improvement. Both participants reacted to the same RDP and could observe both cursors and feedback of immediate and cumulative scores for themselves and the other player (see Methods). Importantly, the task did not enforce a competitive or cooperative context – the individual payoffs were independent. Participants could freely choose whether to use or ignore the perceptual report of the other player.

We pooled all experimental sessions of each participant according to social context (solo vs dyadic, within-subject). Average response lags after a direction change were significantly different in solo and dyadic experiments (Solo: 643 ms ± 79 ms [Mean ± IQR]; Dyadic: 662 ms ± 105 ms; Wilcoxon signed rank test, n = 34, Z = −2.98, p < 0.01). We also found a small but significant improvement in average individual score between solo and dyadic experiments (Solo: 0.236 ± 0.07 [Mean ± IQR]; Dyadic 0.247 ± 0.05; Wilcoxon signed rank test, n = 34 (subjects), Z = 2.31, p < 0.05). Compared to the solo CPR, 68% of participants (23/34) achieved a higher score when co-acting with a partner (**Figure 3A**). On the level of a dyad, the nominal combined score of two co-acting players was higher than that of the same two players working alone in 72% of dyads (Mean difference = 0.023, Wilcoxon signed rank test, n = 50 (dyads), Z= −3.36, p<0.001). Thus, social context seemed to improve overall task outcome as measured by the reward score, although it was not due to an increased number of collected targets (Hit rate Solo: 0.389 ± 0.105 [Mean ± IQR]; Dyadic: 0.391 ± 0.064; Wilcoxon signed rank test, n = 34 (subjects), Z = 0.6582, p = 0.51).

**Figure 3:**
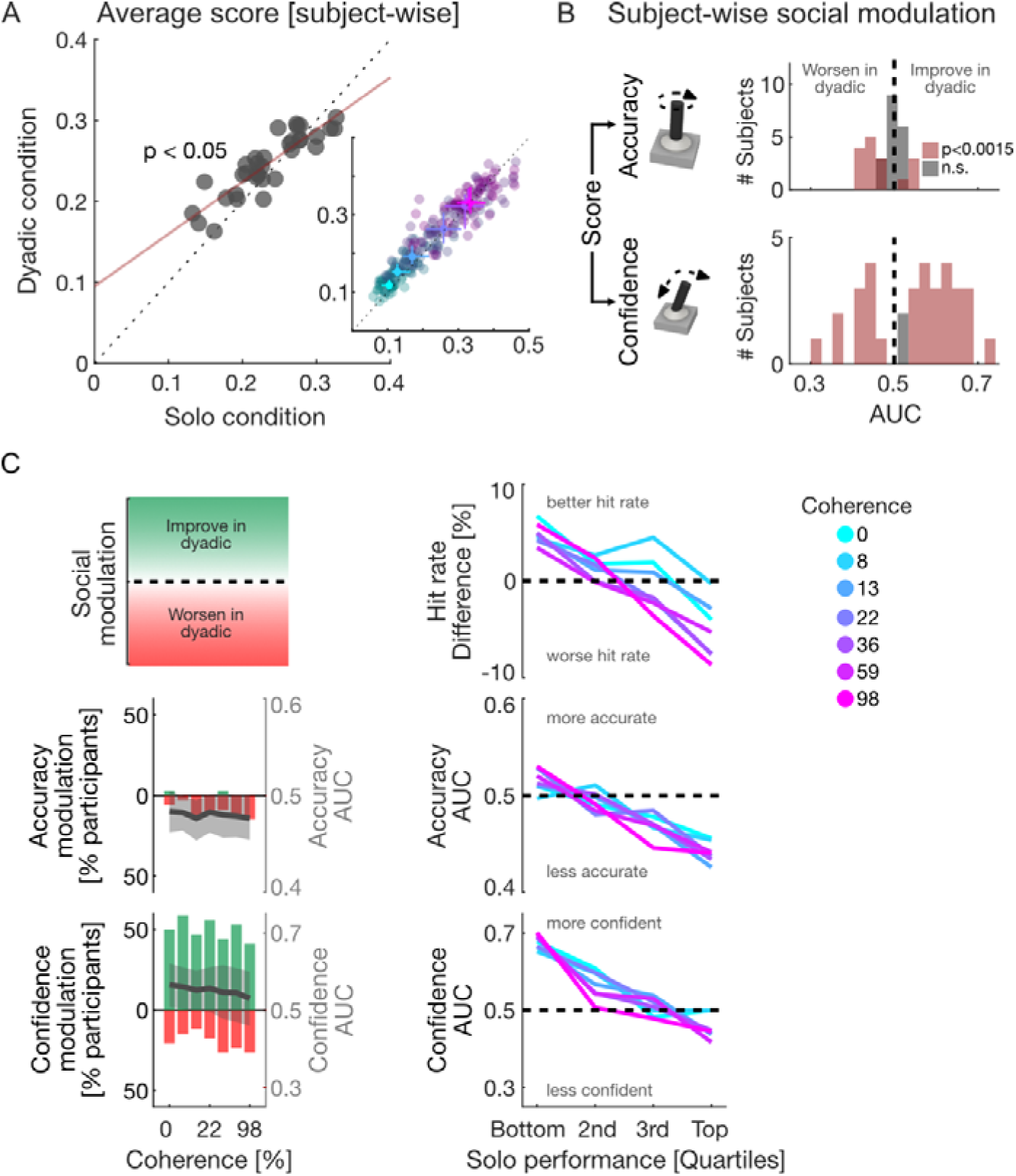
Social modulation: dyadic vs solo. **(A)** Reward score in dyadic vs solo sessions. All solo and (human-human) dyadic sessions were pooled within-subject. Inset: coherence-wise averaging of reward scores. Here and in other panels, stimulus coherence is color-coded. Each subject contributes one data point per stimulus coherence level. The median score across all subjects for each coherence condition is overlaid in brighter color hues. Error bars show 99% confidence intervals of the median in solo and dyadic conditions. **(B)** Social modulation between solo and dyadic experiments, measured as AUC, for state-aligned accuracy (top) and confidence (i.e., tilt, bottom, Wilcoxon rank sum test, Bonferroni-corrected) of individual participants. Coherence was pooled within-subjects. A value of 0.5 corresponds to perfect overlap between solo and dyadic response distributions, 1 and 0 imply perfect separation. See **Supplementary Figure 3A** for average performance in dyadic experiments and **Supplementary Figure 3C** for examples of social modulation and how the AUC captures the its directionality. **(C)** Performance-dependent social modulation. Schematic: social modulation can increase performance (values above the horizontal dashed line, green) or decrease it (values below the horizontal dashed line, red). First column: statistical comparison between joystick accuracy and confidence in solo and dyadic experiments, for each coherence. Sessions were pooled according to experimental condition within-subject. The percentage of participants with significantly different distribution in solo and dyadic sessions is displayed (Wilcoxon rank sum test, Bonferroni-corrected). The directionality of the significant effect in each subject was established with the AUC. Average AUC per coherence level is shown in gray with 99% confidence intervals (shaded background). Second column: average social modulation displayed for different solo performance quartiles, separately for each corresponding performance measure (hit rate, accuracy, confidence). See **Supplementary Figure 3B** for comparison of raw solo joystick responses with social modulation, and **Supplementary Figure 4** for quartile grouping across response dimensions.

Next, we assessed whether changes in perceptual accuracy or confidence were the driving factors behind reward score improvements. To account for the full distributions of accuracy and confidence, we used an AUC analysis to compare performance between solo and dyadic conditions in each subject. Average response accuracy in dyadic experiments did not change for 53% (18/34) of participants and did decline for most of the others (**Figure 3B**; within subjects: Wilcoxon rank-sum test, coherence pooled, number of tests: 34 (subjects), Bonferroni-corrected significance threshold = 0.0015; across subjects: Wilcoxon signed rank test, Z = - 2.25, p < 0.05, Median AUC = 0.485). Thus, the access to reports of others did not improve average CPR competence. However, the social setting altered the confidence reports in 94% (32/34) of participants (**Figure 3B**; within subjects: Wilcoxon rank-sum test, coherence pooled, number of tests: 34 (subjects), Bonferroni-corrected significance threshold = 0.0015; across subjects: Wilcoxon signed rank test, Z = 2.48, p < 0.05, Median AUC = 0.57).

In participants with significantly different confidence in solo versus dyadic conditions, 41% - 59% (min and max across coherence levels) wagered more aggressively on their percept when playing with a partner, while 12% - 26% wagered more conservatively (**Figure 3C**, first column; Wilcoxon rank-sum test, number of tests: 34 (subjects) * 7 (coherence levels) = 238, Bonferroni-corrected significance threshold = 2.1008e-04). Consequently, by signaling more perceptual confidence on their perceptual report in a social setting, most participants achieved a better score.

To understand how social modulation in each participant is influenced by their initial competence or confidence, we sorted the subjects into quartiles based on their solo performance: hit rate, accuracy, and confidence. Generally, participants with low individual performance tended to improve, whereas high-performing ones showed a decline. Initially, less confident participants increased their wagers, while initially more accurate individuals declined in accuracy (**Figure 3C**, second column, see also **Supplementary Figure 4**). Notably, the magnitude of social modulation showed an asymmetry, both with respect to response dimensions and when comparing gains vs losses within each dimension. The least accurate participants improved only slightly, while the most accurate declined more; conversely, the least confident participants gained a lot of confidence, while the most confident lost very little.

These patterns might be explained by convergence between participants, which was shown to impact perceptual accuracy of social decisions (Bang et al., 2017; Mahmoodi et al., 2015; Pescetelli and Yeung, 2022). To directly test the convergence hypothesis, we contrasted the absolute confidence and accuracy differences between the two players in dyadic vs solo settings (**Figure 4A**). Compared to solo experiments, 76% of dyads (38/50) exhibited a smaller difference in confidence when playing together (Wilcoxon signed rank test, n = 50, Z = 3.66, p < 0.001). Similarly, 64% of dyads (32/50) displayed a smaller difference in accuracy (Wilcoxon signed rank test, n = 50, Z = 2.52, p < 0.05). Furthermore, we compared the confidence and the accuracy correlations between the two players, in solo and dyadic contexts. As expected, there was no correlation in solo reports, but the participants’ confidence became significantly correlated when they played together (**Supplementary Figure 5**, Pearson’s correlation, n = 50, Accuracy: Solo: r = −0.04, p = 0.76, Dyadic: r = 0.21, p = 0.14; Confidence: Solo: r = 0.11, p = 0.44, Dyadic: r = 0.54, p < 0.001).

**Figure 4:**
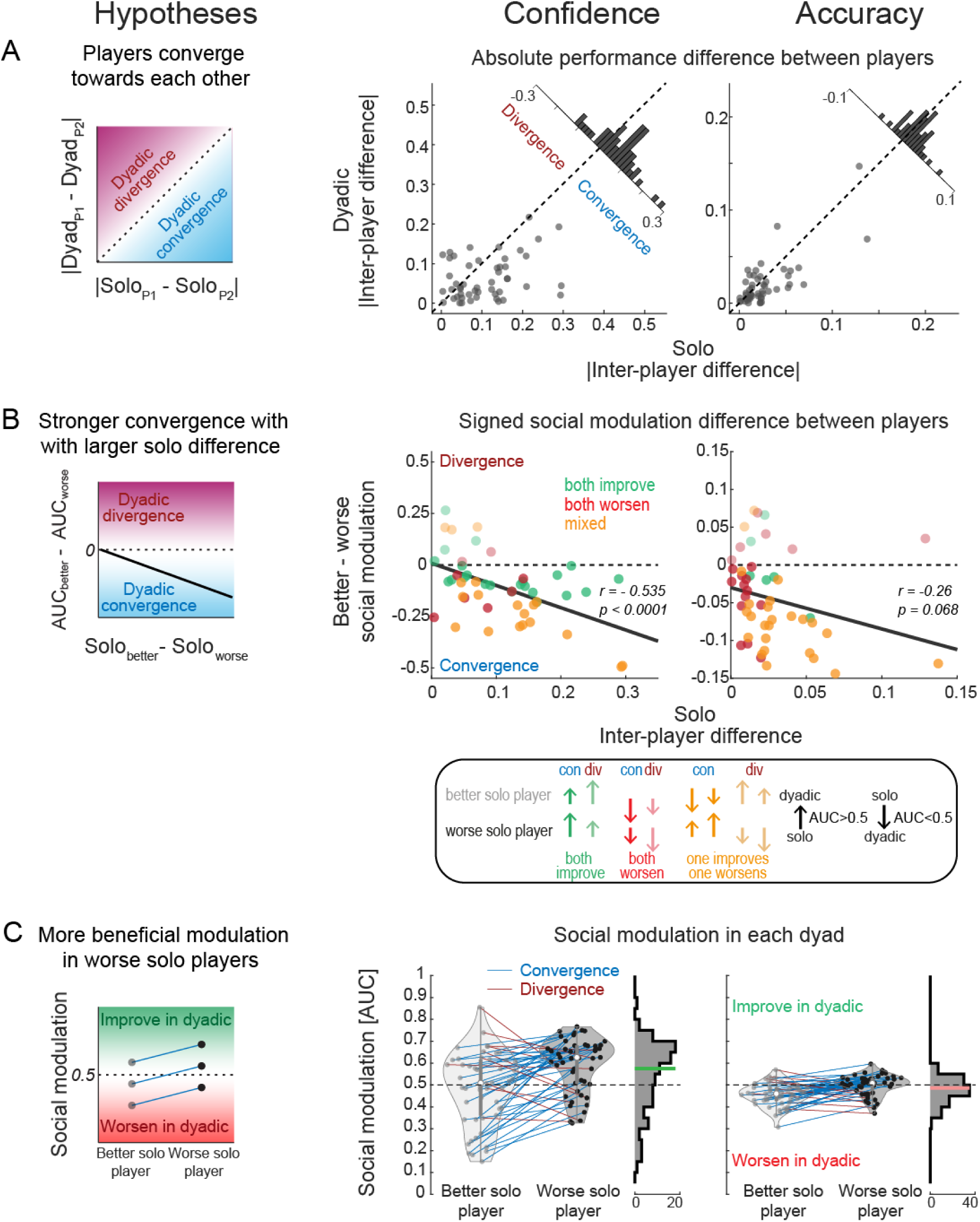
Social modulation in human-human dyads. Left column: schematic depiction of hypothesized effects; middle and right column: actual data for state-aligned confidence and accuracy. **(A)** Absolute difference between partners in solo and dyadic setting for confidence (middle) and accuracy (right). Each dyad is represented by one data point. Dyads show convergence when differences between players in dyadic setting are smaller than differences in solo experiments (see left schematic). **(B)** Social modulation difference between players (better in solo minus worse in solo) as a function of better minus worse inter-player performance difference, for confidence (middle) and accuracy (right). Each dyad is represented by one data point. The solid line illustrates the correlation between solo difference and social modulation difference (Linear regression, n = 50, Confidence: r = −0.535, p < 0.0001; Accuracy: r = −0.26, p = 0.068; Converging dyads only (n = 40): Confidence: r = −0.483, p < 0.01; Accuracy: r = −0.474, p < 0.01). **(C)** Social modulation displayed separately for better and worse solo players. Dyads are connected with a colored line (blue: convergence; red: divergence). Histograms show the overall distribution of social modulations across all participants. Means of the distributions are illustrated with a colored line.

### Performance difference between participants determines social effect

The previous analysis, across subjects, indicated that the participants’ solo performance is an important determinant of how their performance will change in dyadic setting. We next asked whether the amplitude and the direction of social modulation in each participant – summarized with AUC – can be explained by within-dyad differences in solo performance. Intuitively, we hypothesized that a larger difference in solo performance between subjects would lead to a stronger convergence, because little additional information could be derived from observing a similar partner (**Figure 4B**). Theoretically, when considering the better and the worse solo player in a dyad, both could improve in the dyadic setting (AUC > 0.5), both could worsen (AUC < 0.5), or one can improve while the other worsen (here and elsewhere, “worse”/“better” mean less/more confident/accurate, correspondingly).

Depending on the relative strength of the change, this could lead to dyadic convergence or divergence. Confirming the previous result across subjects, 40/50 dyads showed convergence, in both confidence and accuracy. In line with our prediction, we observe stronger convergence between dyadic participants with larger solo difference, indicated by the negative correlation between the better vs worse inter-player AUC difference and the solo performance difference (**Figure 4B**). The initially worse players could exhibit a more beneficial social modulation, relative to the better player, by either improving more or getting less worse (**Figure 4C**, left). Indeed, the initially less confident players improved relative to their counterparts (**Figure 4C**, middle, Wilcoxon signed rank test, n = 50, Z = −4.18, p < 0.001); conversely, less accurate players were barely affected by the social context while their counterparts got worse (**Figure 4C**, right, Wilcoxon signed rank test, n = 50, Z = −4.35, p < 0.001). This asymmetric pattern resulted in an overall positive shift for confidence and slight overall negative shift for accuracy (**Figure 4C**, histograms, Wilcoxon signed rank test against 0.5, n = 100; Confidence: Median AUC = 0.5752, Z = 2.80, p < 0.01; Accuracy: Median AUC = 0.486, Z = −3.55, p < 0.001). Importantly, even though we analyzed the confidence and the accuracy separately, there was a strong overlap between these two measures within a dyad: the initially more confident player was also the more accurate one in 43/50 (86%) dyads. This allows interpreting the plots in **Figure 4C** across panels: e.g. not only less confident but also less *accurate* participants tend to gain *confidence* (see also **Supplementary Figure 4**).

In contrast to earlier studies that found more successful perceptual decision-making in social settings when perceptual sensitivities or confidence match between participants (Bahrami et al., 2010; Pescetelli and Yeung, 2022), our data does not reveal systematic “dyadic benefits” for dyads with similar perceptual accuracy or confidence. Instead, there was a positive (but not significant) relationship between the average social modulation within a pair and solo difference between players (**Supplementary Figure 6**).

### Perceptual accuracy improves with reliable social signaling

As expected, the quality of the solo perceptual report declined in a comparable fashion across participants for low stimulus coherence (**Figure 2**). We wondered if this perceptual breakdown led to the relatively small accuracy modulation by the social context we described above (**Figure 3** and **Figure 4**). Based on earlier work on Bayesian integration and social conformity (De Martino et al., 2017; Germar et al., 2016; Khalvati et al., 2021; Park et al., 2017), we expected that integrating information from a partner will be weighted by their reports’ accuracy reliability. We hypothesized that participants would integrate more social evidence when it was reliably accurate, regardless of the stimulus noise. Furthermore, we asked whether incorporating social signals into human decision-making requires graded, accuracy-depended confidence signaling by others.

To that end, we developed a computer player, that was programed to accurately represent the nominal RDP direction (± Gaussian noise; note that even 0% coherence had a correct “nominal” direction), with a fixed “confidence” of 0.5 (± Gaussian noise in a.u.) across all coherence levels at all times (**Supplementary Figure 7**). In such human-computer (HC) dyads, the computer player was physically impersonated by one of the experimenters who pretended to be the partner. Thus, participants believed that they played the game with another human. The computer player was set up to report motion direction with a constant, human-like latency (508 ms ± Gaussian noise). The computer response was not affected by the cursor of the human participants, resulting in a situation where the social signals might only unilaterally affect the human player. Crucially, unlike real human partners, the computer player did not provide useful information regarding its tracking confidence. With this condition we aimed to evaluate whether human participants would integrate social cues about the motion direction into their own reports, while the partner’s confidence report was uninformative. We nevertheless expected a response accuracy improvement, especially when sensory evidence became degraded at low coherences. Furthermore, we hypothesized that the reliable nature of the computer partner would result in riskier, more eccentric joystick placement in human players.

By accurately representing the stimulus direction, the computer player triggered profound behavioral effects (**Supplementary Table 2** to **Supplementary Table 4**). In this setting, participants collected more targets with a higher score (**Figure 5A**). Compared to human-human dyads, 63% of participants (21/33) improved their accuracy (and only 3 participants became worse) across all coherence levels (**Figure 5B-C**, within-subjects: Wilcoxon rank-sum test, coherence pooled, number of tests = 33 (subjects), Bonferroni-corrected significance threshold = 0.0015; across subjects: Wilcoxon signed rank test, Median AUC = 0.56, Z = 3.87, p < 0.001; see **Supplementary Figure 8** for comparison to solo behavior). In particular, 55% of participants (18/33) showed a significant accuracy boost at 0% coherence (Wilcoxon rank-sum test, number of tests: 33 (subjects) * 7 (coherence levels) = 231, Bonferroni-corrected significance threshold = 2.1645e-04, Median AUC across subjects at 0% coherence = 0.66). Thus, participants integrated reliable sensory-social direction cues to improve their task performance, especially when stimulus was ambiguous, suggesting a unilateral convergence towards the computer player.

**Figure 5:**
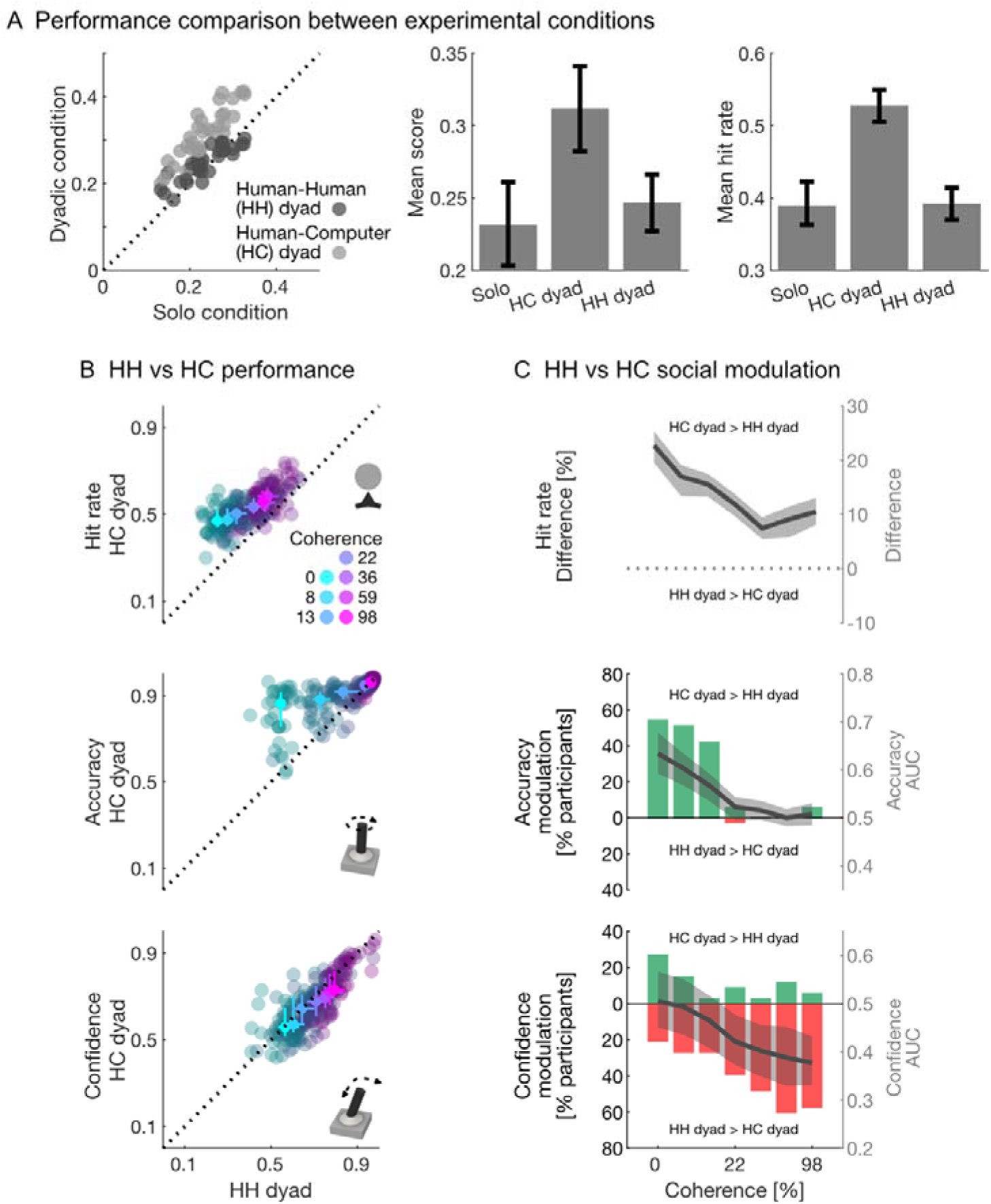
Comparison of social modulation in human-human (HH) dyads and human-computer (HC) dyads. **(A)** Left: average subject-wise score in the two dyadic experiments compared to the score of the same participant in solo experiments. Average score (middle) and hit rate (right) of the population in different experimental conditions. Error bars correspond to the 99% confidence intervals of the mean. **(B)** Comparison of average hit rate (top), accuracy (middle) and confidence (bottom) for each participant and each stimulus coherence (color-coded) in human-human and human-computer dyads. Individual data (subject-wise, averaged across several HH sessions for each subject) are shown in darker hue. Medians are overlaid for each coherence condition with bright colors. Error bars show 99% confidence intervals of the median. **(C)** Statistical comparison of accuracy and confidence in HH and HC dyadic experiments, for each coherence condition. Sessions were pooled according to experimental condition within-subject. The percentage of participants with significantly different distribution in HH vs HC dyadic sessions is displayed (Wilcoxon rank sum test, Bonferroni-corrected). The directionality of the significant effect in each subject was established with the AUC. Average AUC per coherence level is shown in gray with 99% confidence intervals (shaded background). For hit rates, the average difference is displayed.

Despite more accurate task performance, most participants did not improve in the confidence dimension (**Figure 5B-C**). In fact, compared to an interaction with a real human counterpart, 64% of participants showed less confidence while only 18% improved when playing with the computer player (within-subjects: Wilcoxon rank-sum test, coherence pooled, number of tests = 33 (subjects), Bonferroni-corrected significance threshold = 0.0015, across subjects: Wilcoxon signed rank test, Median AUC = 0.45, Z = −3.15, p < 0.01). Subjects were particularly affected when the task was easy (98% coherence, Median AUC across subjects = 0.36). This too seems to suggest confidence convergence towards the relatively invariant low confidence computer player, even with otherwise reliably accurate direction signaling.

### Social modulation of confidence and accuracy co-varies

So far, we analyzed the social modulation of perceptual confidence and accuracy independently. Here we investigate the link between the two response dimensions. In both dyadic conditions, subject-wise social modulation of perceptual confidence correlated with the change in accuracy (**Figure 6**, HH: Pearson’s correlation, n = 100, r = 0.76, p < 0.001; HC: n = 33, r = 0.56, p < 0.001), suggesting that the gain or the loss in accuracy leads to reappraisal of confidence. Note however that a substantial fraction of participants had an incongruent social modulation, showing *less* accuracy but *more* confidence (**Figure 6**, HH, upper left quadrant).

**Figure 6:**
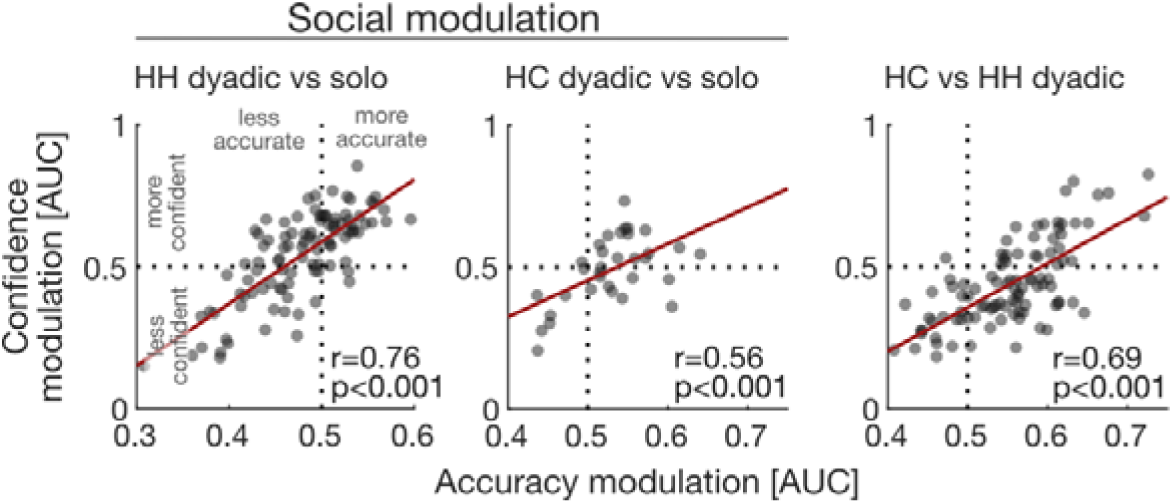
Relationship between social modulation of accuracy and confidence in each participant. Each dot represents one player in a dyad. Left: Social modulation in human-human (HH) dyadic condition vs. solo (n = 100); Middle: Human-Computer (HC) dyadic condition vs solo (n = 33); Right: Correlation of confidence vs accuracy difference between two dyadic conditions (HC vs HH, n = 98). Values above 0.5 on each axis correspond to positive social modulation: increased accuracy or confidence in the “first condition” (e.g. HH) compared to the “second condition” (e.g. solo), and vice versa.

Despite the positive covariation between social modulation of accuracy and confidence, the human–computer experiment demonstrated that social evidence integration does not require metacognitively sensitive, graded confidence signaling by the partner. However, the apparent dissociation between the *average* improvement of accuracy and the *average* decline of confidence is still grounded in a lawful relationship between two response dimensions. Players who gained substantially more accuracy tended to show confidence increase, or at least less confidence decrease, compared to players who did not change much in accuracy (**Figure 6**, HC vs HH: n = 98, r = 0.62, p < 0.001). To conclude, both, individually-varied, coherence-dependent human reports, and reliably accurate direction reports coupled with uninformative confidence expression by the simulated partner, influenced human perceptual decisions and resulted in converging, socially conforming behavior.

## Discussion

In this study, we assessed continuous human perceptual decision-making in individual and social settings, with a newly developed paradigm, where subjects wager on the correctness of their motion percept in real-time. Overall, during dyadic co-action, we find higher perceptual confidence but no gain in accuracy. We demonstrate that asymmetric convergence underlies this net effect, with the magnitude and directionality of the social modulation depending on the difference between competence and confidence of the partners.

In contrast to the increase in confidence in the dyadic condition we have observed, previous social perceptual decision studies did not report overall rise in confidence (Bang et al., 2017; Pescetelli et al., 2016; Pescetelli and Yeung, 2022). At the same time, some earlier work demonstrated gains in dyadic competence – i.e. when the group decision-making outperforms even the better of two partners – after partners exchanged their individual confidence and made a joint choice (Bahrami et al., 2010; Pescetelli et al., 2016). Likewise, competence increases even during dyadic co-action – without interdependent payoff or joint decision – especially for participants with similar confidence (Pescetelli and Yeung, 2022). But despite presenting subjective confidence continuously and saliently as part of the perceptual report, our task did not elicit overall – across all participants – improvement in accuracy or hit rate. The resulting score (which combines accuracy, hit rate, and confidence), however, did improve in the dyadic setting, reflecting a dyadic benefit driven by an overall increase in confidence. This indicates that real-time social feedback boosts confidence in one’s perception, even in the absence of a corresponding enhancement in competence. Such a disconnect between the effects on confidence (which can be construed as subjectively perceived competence) and actual competence is reminiscent of previous work on metacognitive biases in individuals (Kruger and Dunning, 1999): many less competent participants showed socially-induced increase in confidence without an associated increase – or even with a decrease – in accuracy.

These effects were driven by a mutual convergence between dyadic partners along both the perceptual accuracy and the confidence dimensions. For each participant, the direction and the magnitude of social modulation was determined by the difference in the initial performance of the two players, with larger solo differences between participants resulting in larger difference in social modulation between the partners. Critically, dyadic convergence affects both partners, but often asymmetrically. For instance, the more confident players on average show very little change in confidence but they strongly boost the confidence of their partners. Dyadic confidence convergence (Esmaily et al., 2023; Pescetelli and Yeung, 2022) and confidence matching (Bang et al., 2017) have been described before. An elegant collective decision study showed that an “equality bias” – assigning similar weights to both participants regardless of their respective competence – might underlie mutual convergence (Mahmoodi et al., 2015). Interestingly, in their study more competent dyad members were better at weighting their partner’s options compared to less competent ones. In contrast, in our experiment more competent players lose more accuracy than is gained by their partners. Furthermore, unlike the previous work, where similar confidence (Pescetelli and Yeung, 2022) or perceptual sensitivity (Bahrami et al., 2012a, 2010; Baumgart et al., 2020) correlated with higher dyadic benefit, we do not find systematic dyadic competence benefits (in our case, nominal “average accuracy” within a dyad) for participants with similar task competence or confidence. We speculate that the mode of social feedback and interaction might underlie these differences. Explicit communication and a joint decision – made together or by one player (Bahrami et al., 2012a, 2010; Baumgart et al., 2020), or periods of metacognitive introspection in which prior individual decisions are evaluated (Pescetelli and Yeung, 2022), might have elicited competence improvements in those studies.

The control experiment with a reliably accurate, simulated dyadic “partner” who exhibited a stable but fairly low level of confidence irrespective of the task difficulty elicited vastly improved accuracy and hit rate, especially when sensory information was ambiguous. At the same time, participants’ confidence reports became more conservative when playing with such “conservative” partner, because participants gravitated towards the report of the simulated partner. Improvements in competence coupled with declining confidence further support dyadic convergence where the direction can be dissociated along the two dimensions. Thus, instead of using the reliably accurate information provided by the computer player to be more accurate *and* more confident, convergence interfered with fully maximizing the reward score. High-accuracy, low-confidence simulated partners have been recently shown to elicit more conservative confidence reports during binary dyadic decision-making (Esmaily et al., 2023). Beyond these results, our experiments demonstrate that humans do not require sensible confidence expression to recognize and utilize differences in task competence. Our findings indicate a possible dissociation between the accuracy and confidence alignment. At the same time, a positive covariation of accuracy and confidence modulation by the dyadic context suggests a retention of metacognitive sensitivity under social influence. Thus, performance history and temporal reliability might be important factors in addition to explicitly signaled confidence, especially when these information streams are not congruent.

Systematic changes towards group consensus (‘social conformity’) have been shown to bias decision-making towards majority choices (De Martino et al., 2017; Germar et al., 2016; Park et al., 2017; Toelch and Dolan, 2015). The dyadic convergence we and others observe might be the basis for social conformity in larger group settings. These findings resonate with the work on factual knowledge estimation by Lorenz and colleagues, who showed that social influence can reduce the diversity of opinions, leading to increased confidence without improved group accuracy (Lorenz et al., 2011). Reinterpreting these results, Farrell argued that information sharing can enhance individual performance and confidence calibration, even if group-level diversity diminishes. This statistical “shrinkage” effect may benefit individuals by reducing extreme errors. Thus, while social influence may undermine the classical “wisdom of crowds” at the aggregate level, it can simultaneously improve individual outcomes and self-assessment (Farrell, 2011; Lorenz et al., 2011; Rauhut et al., 2011). Our results suggest that real-time social feedback may provide similar individual-level benefits, even if it fails to enhance collective perceptual competence.

These nuanced relationships between individual and joint – often only nominally “joint” – success underscore the importance of the actual social context. Similar to many social influence studies such as above, in our dyadic condition participants were not instructed to cooperate or compete with one another, nor incentivized to perform better as a group. Instead, they co-acted under independent reward contingencies. This difference to joint decision studies (Bahrami et al., 2012b, 2012a, 2010; Baumgart et al., 2020; Esmaily et al., 2023; Mahmoodi et al., 2015; Pescetelli et al., 2016) is crucial for interpreting our results, since participants could ignore the overt behavior of the other player. Therefore, any social modulation or correlated behavior observed in our experiment can be attributed to a spontaneous, self-regulated process. Supporting a prior perceptual decision co-action study (Pescetelli and Yeung, 2022), we interpret our findings as evidence that in social situations people spontaneously and opportunistically integrate the judgment of others into their own decisions, even when social interaction is not incentivized or enforced. In line with this argument, humans seem to naturally follow gaze signals and choice preferences of others, suggesting the utilization of others’ thoughts and intentions (Bayliss et al., 2007; Madipakkam et al., 2019; Mitsuda and Masaki, 2018). Furthermore, human co-action seems to result in attentional attraction or withdrawal in some dyads (Dosso et al., 2018). As next step, it would be very interesting to test whether face-to-face interactions through the transparent shared visual display will induce even stronger social effects compared to separate experimental booths (Isbaner et al., 2025; Lewen et al., 2025; Moeller et al., 2023).

The advent of new techniques such as time-continuous decision-making (Bonnen et al., 2015; Huk et al., 2018; Noel et al., 2023, 2022) and hyperscanning (Babiloni and Astolfi, 2014; Czeszumski et al., 2020) allows to ask how evolving decisional variables are represented in neural circuitry underlying flexible behaviors. This is an important step beyond the traditional approach based on discrete, trial-based decisions. Adapting this approach, we demonstrate real-time, dynamic influence of social information on human perceptual decisions. Studies investigating the neuronal correlates of similar perceptual decisions have demonstrated faster and more accurate behavioral responses when the sensory evidence resulted in earlier and more reliable neuronal changes (Fan et al., 2018; Gold and Stocker, 2017; Kiani and Shadlen, 2009). It has also been shown that microstimulation- and optogenetically-elicited inputs can be integrated into perceptual decisions (Fetsch et al., 2018, 2014; Salzman et al., 1990). Along these lines, we propose that reliably accurate real-time social information is multiplexed with sensory signals, possibly resulting in enhanced encoding already in cortical neurons representing relevant sensory dimensions.

In summary, our novel CPR task is a powerful new tool for studying the dynamics of decision and confidence formation, in individual and social settings. We show that the presence of a co-acting social partner adaptively changes continuous and graded perceptual decisions, resulting in mutual but asymmetric convergence and often a net dyadic benefit in confidence and the reward score. This is particularly apparent in a confidence boost of a less confident – and often less accurate – dyadic partner. On the other hand, more accurate partners on average lose accuracy. The lawful relationships between confidence and competence modulations demonstrate the importance of concurrently considering these two measures, both within each participant and across interacting partners. These results advance our understanding of how humans evaluate and incorporate social information, especially in real-time decision-making situations that do not permit careful but slow deliberations.

## Methods

### Study design and participants

Data were recorded from 38 human participants (Median age: 26.17 years, IQR 4.01; 13 of which with corrected vision) between January 2022 and August 2023. Prior to the experimental sessions, each participant was trained on two occasions. During the experimental phase, participants played three variations of the experimental paradigm: alone (solo), with a human player (Human-Human dyad), and with a computer player (Human-Computer dyad). The experimental order was mixed and largely determined by the availability of participants (Supplementary Figure 1A). Most participants in this study originated from central Europe or South Asia. All procedures performed in this study were approved by the Ethics Board of the University of Göttingen (Application 171/292).

### Experimental setup

Participants sat in separate experimental booths with identical hardware. They were instructed to rest their head on a chinrest, placed 57 cm away from the screen (Asus XG27AQ, 27” LCD). A single-stick joystick (adapted analog multifunctional joystick (Sasse), Resolution: 10-bit, 100 Hz) was anchored to an adjustable platform placed in front of the participants, at a height of 75 cm from the floor. The joystick was calibrated before data acquisition to ensure comparable readouts. Screens were calibrated to be isoluminant. Two speakers (Behringer MS16), one for each setup were used to deliver auditory feedback at 70 dB SPL.

The experimental paradigm was programmed in MWorks (Version 0.10 – https://mworks.github.io). Two iMac Pro computers (Apple, MacOS Mojave 10.14.6) served as independent servers for each setup booth. These computers were controlled by an iMac Pro (Apple, MacOS Mojave 10.14.6). Custom-made plugins for MWorks were used to generate and display the stimuli, to handle the data acquisition from the joystick (10 ms sampling rate), and to incorporate all data from both servers into a single data file.

### Continuous perceptual report (CPR) game

Participants were instructed to maximize monetary outcome in a motion tracking game. In this game, subjects watched a frequently changing random dot pattern on the screen and used a joystick (Szul et al., 2020) to indicate their current motion direction perception. The joystick controlled an arc-shaped response cursor on the screen (partial circle with fixed eccentricity, Solo: 2 degree of visual angle (‘dva’) radius from the center of the screen; Dyadic: 1.8 dva & 2 dva radius). The angular direction of the joystick was linked to the cursor’s polar center position. In addition, the joystick tilt was permanently coupled to the cursor’s width (see below, 13 – 180 degrees). This resulted a continuous representation of the joystick position along its two axes. By moving the joystick, participants could rotate and shape the cursor. At unpredictable times (1% probability every 10 ms), a small white disc (‘reward target’, diameter: 0.5 dva) appeared on the screen for a duration of 50 ms at 2.5 dva eccentricity, congruently with the motion direction of the stimulus. Whenever a target appeared in line with the cursor, the target was considered collected (‘hit’) and the score of the participant increased. To help participants performing such alignment to the best of their perceptual abilities, a small triangular reference point was added to the center point of the cursor. Throughout the experiment, participants were required to maintain gaze fixation on a central fixation cross (2.5 dva radius tolerance window) or the cursor would disappear and no targets could be collected until fixation was resumed.

In the solo experiments, the cursor was always red. In dyadic conditions, the two cursors, present on screen simultaneously, were red and green (isoluminant at 17.5 cd/m^2^ ± 1 cd/m^2^). During dyadic experiments, the position of the two cursors switched between stimulus cycles, with the red cursor always starting above, but not overlapping the green cursor. Each cursor color was permanently associated with one of the two experimental booths. After the mid-session break, participants switched booths, contributing an equal amount of data for each setup (600 reward targets, ∼20 min, up to 17 stimulus cycles). Players initiated new stimulus cycles with a joystick movement. Each stimulus cycle could last up to 75 seconds, during which the RDP’s motion direction and coherence changed at pseudorandomized intervals, resulting in the presentation of 30 stimulus states per cycle.

### Random dot pattern (RDP)

We used a circular RDP (8 dva radius) with white dots on a black background. Each dot had a diameter of 0.1 dva, moved with 8 dva/s and had a lifetime of 25 frames (208 ms). The overall dot density was 2.5 dots/dva. The stimulus patch was centrally located on the screen. The central part of the stimulus (5 dva diameter) was blacked out. In this area we presented the fixation cross and the response arc. The RDP motion direction was randomly seeded and set to change instantly by either 15 deg, 45 deg, 90 deg or 135 deg after a pseudo-randomized time interval of 1250 ms to 2500 ms. Whether the signal moved in clockwise or counterclockwise direction was random. Only signal dots altered their direction. The dot coherence changed pseudo-randomly after 10 RDP direction changes to the coherence level that was presented least. Seven coherence conditions were tested: 0, 8, 13, 22, 36, 59, 98%.

### Gaze control

Participants were required to maintain gaze fixation at the center of the screen throughout each stimulus cycle. We used a white cross (0.3 dva diameter) as anchor point for the participants’ gaze. The diameter of the fixation window was set to 5 dva. An eye tracker (SR research, EyeLink 1000 Plus) was used to control gaze position in real-time. If the gaze position left the fixation window for more than 300 ms, the player’s arc would disappear from the screen, preventing target collection. In addition to this, an increase of the fixation cross’ size, together with a change in color (white to red), signaled to the participants that fixation was broken. As soon as the gaze entered the fixation window again, visual parameters were reset to the original values and the arc would reappear, allowing the player to continue target collection.

### Reward score

Participants were incentivized to maximize their monetary outcome by collecting as many targets as possible with the highest possible score. The minimum polar distance between the arc’s center position and the nominal direction (target center, ‘accuracy’) as well as the angular width of the response cursor (‘joystick tilt’) at the moment of collection were taken into account when calculating the score:

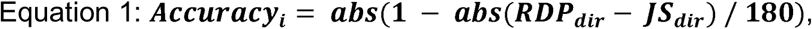

where *RDP_dir_*refers to the direction of the random dot pattern and *JS_dir_* refers to the direction of the joystick at sample *i*.

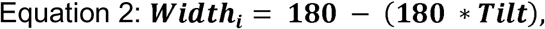

where *tilt* varies between 0 and 1 for minimal and maximal radial joystick positions, respectively.

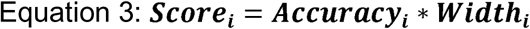

Thus, narrower and more accurately placed cursors caused higher reward scores.

### Feedback signals

Various feedback signals were provided throughout the experiment to inform participants about their short- and long-term performance. All feedback signals were mutually visible.

### Immediate feedback

Immediately after a target was collected, visual and acoustic signals were provided simultaneously. The auditory feedback consisted of a 200 ms long sinusoidal pure tone at a frequency determined by the score. Each tone corresponded to a reward range of 12.5%, with lower pitch corresponding to low reward score. We used 8 notes from the C5 major scale (523, 587, 659, 698, 784, 880, 988, 1047 Hz). Sounds were on- and off-ramped using a 50 ms Hanning window. No sound feedback was given for missed targets. In solo experiments, the visual feedback consisted of a 2 dva wide circle, filled in proportion to the score with the same color as the arc’s player. The circle was presented in the center of the screen, behind the fixation cross for 150 ms. In dyadic conditions, the visual feedback consisted of half a disc for each player and both color-coded halves were mutually visible.

### Short-term feedback

During each stimulus cycle, a running average of the reward score was displayed for each player with a 0.9 dva wide, color-coded ring around the circumference of the RDP (18.2 dva and 19.4 dva diameter). After every target presentation, the filled portion of the ring updated. To avoid spatial biasing, the polar zero position of the ring changed randomly with every stimulus cycle.

### Long-term feedback

Cumulative visual feedback was provided after each stimulus cycle (during the inter-cycle intervals) for 2000 ms. It displayed the total reward score accumulated across all cycles as a colored bar graph located at the center of the screen. A grey bar (2 dva wide, max height: 10 dva) indicated the maximal possible cumulative score after each cycle. A colored bar next to it (same dimensions) showed how much was collected by the player so far. In dyadic experiments, red and green bars would be shown on either side of the grey bar. In solo experiments, a red arc was shown to the left of the grey bar. The configuration of the visual stimuli and task parameters is illustrated in **Figure 1** and as a supplementary video file **Supplementary Video 1**.

### Statistical analysis

Performance metrics, based on raw joystick tilt and accuracy, were extracted and averaged in time windows of 30 frames (∼250 ms) that were either target- or state-aligned. Target-alignment refers to time windows prior to the first reward target presentation of each stimulus state. Target-aligned data were only considered if the first target appeared at least 1000 ms after the direction change. This analysis approach was chosen to (i) allow adequate time for an intentional response and (ii) to avoid the prediction of motion direction based on earlier target locations. State-alignment refers to a 30 frames time window before a nominal motion direction change of the stimulus (end of the current stimulus state). Stimulus states in which fixation breaks exceeded 10% of the state duration were excluded from the performance analyses.

For population analyses across subjects, joystick response parameters (tilt and accuracy) were first averaged within-subject. Bootstrapped 95% confidence intervals were estimated with 1000 repetitions (Matlab: bootci). For analyses within each subject or a dyad, differences of accuracy and tilt between experimental conditions (solo vs dyadic) were tested with a two-sided Wilcoxon signed rank test (Matlab: signrank, for paired samples) and a two-sided paired Wilcoxon rank sum test (Matlab: ranksum). Bonferroni correction was applied for multiple testing. Whether or not coherence was pooled is indicated for each test. Within-subject effect size and direction was estimated with the area under the receiver operating characteristic (‘AUC’, Matlab: perfcurve). AUC values of 0.5 indicated similar distributions. AUC of 0 and 1 suggested perfectly separated distributions. Directionality of social modulation was inferred by AUC change (larger vs smaller than 0.5). Correlation coefficients were calculated with a Pearson correlation (Matlab: corrcoef). Average response lags between stimulus and response were estimated with the maximum cross-correlation coefficient (Matlab: xcorr). Social modulation differences between dyadic players were compared to the baseline modulation of shuffled dyadic partners (Matlab: randperm). We fitted three Generalized Linear Mixed Models (GLMM; Baayen, 2008) to the raw joystick responses parameters, which differed in their response variable and the size of the data set analyzed but had identical fixed effects structures and largely identical random effects structures. We fitted one model for the probability of a target hit (model 1a), joystick tilt (model 1b), and joystick accuracy (model 1c) as the response. All three aimed at estimating the extent to which the respective response variable was affected by the fixed effects of experimental condition (solo or human-computer dyad), random dot pattern coherence, stimulus duration, stimulus number, block number, and day number. We hypothesized that the effect of coherence depended on the condition, thus, we included the interaction between these two predictors into the fixed affects part of the model. To avoid pseudo-replication and account for the possibility that the response was influenced by several layers of non-independence, we included three random intercepts effects, namely those of the ID of the participant, the ID of test day (nested in participant; thereafter ‘day ID’), and the ID of the block (nested in participant and day; thereafter ‘block ID’). The reason for including the latter two was that it could be reasonably assumed that the performance of participants varied between test days and also between blocks tested on the same day. To avoid an ‘overconfident model’ and keep type I error rate at the nominal level of 0.05 we included all theoretically random slopes (Barr et al., 2013; Schielzeth and Forstmeier, 2009). These were those of condition, coherence, their interaction, stimulus duration, stimulus number, block number, and day number within participant, coherence, stimulus duration, stimulus number, and block number within day ID, and finally coherence, stimulus duration, and stimulus number within block ID. Originally we also included estimates of the correlations among random intercepts and slopes into each model, but do to convergence and identifiability problems (recognizable by absolute correlation parameters being close to 1; Matuschek et al., 2017) we had to exclude all or several of these estimates from the full models (see **Supplementary Table 1** for detailed information).

For each model we conducted a full-null model comparison which aims at avoiding ‘cryptic multiple testing’ and keeping the type one error rate at the nominal level of 0.05 (Forstmeier and Schielzeth, 2011). As we had a genuine interest in all predictors present in the fixed effects part of each model the null models comprised only the intercept in the fixed effect’s part but were otherwise identical to the respective full model. This full-null model comparison utilized a likelihood ratio test (Dobson, 2001). Tests of individual effects were also based on likelihood ratio test, comparing a full model with an each in a set of reduced models which lacked fixed effects one at a time.

### Model implementation

We fitted all models in R (version 4.3.2; R Core Team, 2023). In model 1a we included the response as a two columns matrix with the number of targets hit and not hit in the first and second column respectively (Baayen, 2008). The model was fitted with a binomial error structure and logit link function (McCullagh and Nelder, 1989). In essence, such models model the proportion of targets hit. We are aware that in principle one would need an ‘observation level random effect’ which would link the number of targets hit and not hit in a given stage. However, in a relatively large proportion of stages (19.7%) there was only a single target that appeared and in the majority of stages (47.0%) only two targets appeared, making it unlikely that a respective random effect can be fitted successfully.

For models 1b and 1c, we fitted with a beta error distribution and logit link function (Bolker, 2008). Models fitted with a beta 1 error distribution cannot cope with values in the response being exactly 0 or 1. Hence, when such values were present in a given response variable we transformed then as suggested by Smithson & Verkuilen, 2006. Model 1a was fitted using the function glmer of the package lme4 (version 1.134; Bates et al., 2015), and models 1b and 1c were fitted using the function glmmTMB of the equally named package (version 1.1.8; Brooks et al., 2017). We determined model stability by dropping levels of the random effects factors, one at a time, fitting the full model to each of the subsets, and finally comparing the range of fixed effects estimates obtained from the subsets with those obtained from the model fitted on the respective full data set. This revealed all models to be of good stability. We estimated 95% confidence limits of model estimates and fitted values by means of parametric bootstraps (N=1000 bootstraps; function bootMer of the package lme4 for model 1 and function simulate of package glmmTMB for models the response was overdispersed (maximum dispersion parameter: 1.0).

## Glossary

### Dyadic condition

Real-time co-action of two players, each having instant access to the partner’s report but without payoff interdependence. One player is always a human participant; the other can be either a real human participant (HH dyad) or a simulated computer agent (HC dyad), which was impersonated by the experimenter. *Hit rate*: Fraction of successfully collected reward targets. Success (hit) was achieved when the joystick-controlled cursor overlapped with the reward target at the time of its appearance.

### Joystick accuracy

Inverse normalized tracking error of the joystick. A value of 1 corresponds to a joystick direction response in nominal stimulus direction. A value of 0 indicates a response which is 180 degrees off the nominal stimulus direction. Used as measure for task competence.

### Joystick tilt

Refers to the normalized radial displacement of the joystick from the center position. A value of 1 indicates maximum displacement from the joysticks center position. Used as proxy measure for perceptual confidence.

### Reward targets

Small stimuli which were briefly presented at pseudo-random times in nominal stimulus direction. Subjects were instructed to collect these targets with their joystick-controlled cursor, to obtain rewards.

### Reward score

Normalized monetary payoff. Calculated as the product of normalized accuracy and tilt at time of reward target presentation for hits, and is always zero for misses.

### Stimulus state

A pseudorandomized epoch of time with constant nominal direction and coherence of the random dot pattern.

## Acknowledgements

This publication was funded by the Deutsche Forschungsgemeinschaft (DFG, German Research Foundation) - Project-ID 454648639 - SFB 1528 - Cognition of Interaction, subproject A01, and the Leibniz Collaborative Excellence grant K265/2019 “Neurophysiological mechanisms of primate interactions in dynamic sensorimotor settings” (PRIMAINT) and the European Union (ERC Starting Grant, NEUROGROUP, 101041799). Views and opinions expressed are however those of the authors only and do not necessarily reflect those of the European Union or the European Research Council Executive Agency. Neither the European Union nor the granting authority can be held responsible for them. We thank Fred Wolf for useful discussions.

## Author contribution

Conceptualization: F.S., A.C., A.G., I.K., S.T.; Methodology: F.S., A.C., S.T.; Investigation: F.S.; Analysis: F.S., R.M.; Software: F.S.; Visualization: F.S., I.K.; Writing – Original Draft: F.S., A.C., I.K.; Writing – Review & Editing: all authors; Funding Acquisition: A.G, I.K., S.T.; Resources: S.T.; Supervision: I.K, S.T.

## Lead contact

Further information and requests for resources should be directed to and will be fulfilled by the lead contact, Felix Schneider.

## Data and code availability

The dataset and MATLAB code generated during this study is available at GitHub (https://github.com/SocCog-Team/CPR/tree/main/Publications/2024_perceptual_confidence).

## Supplementary Figures

**Supplementary Figure 1.**
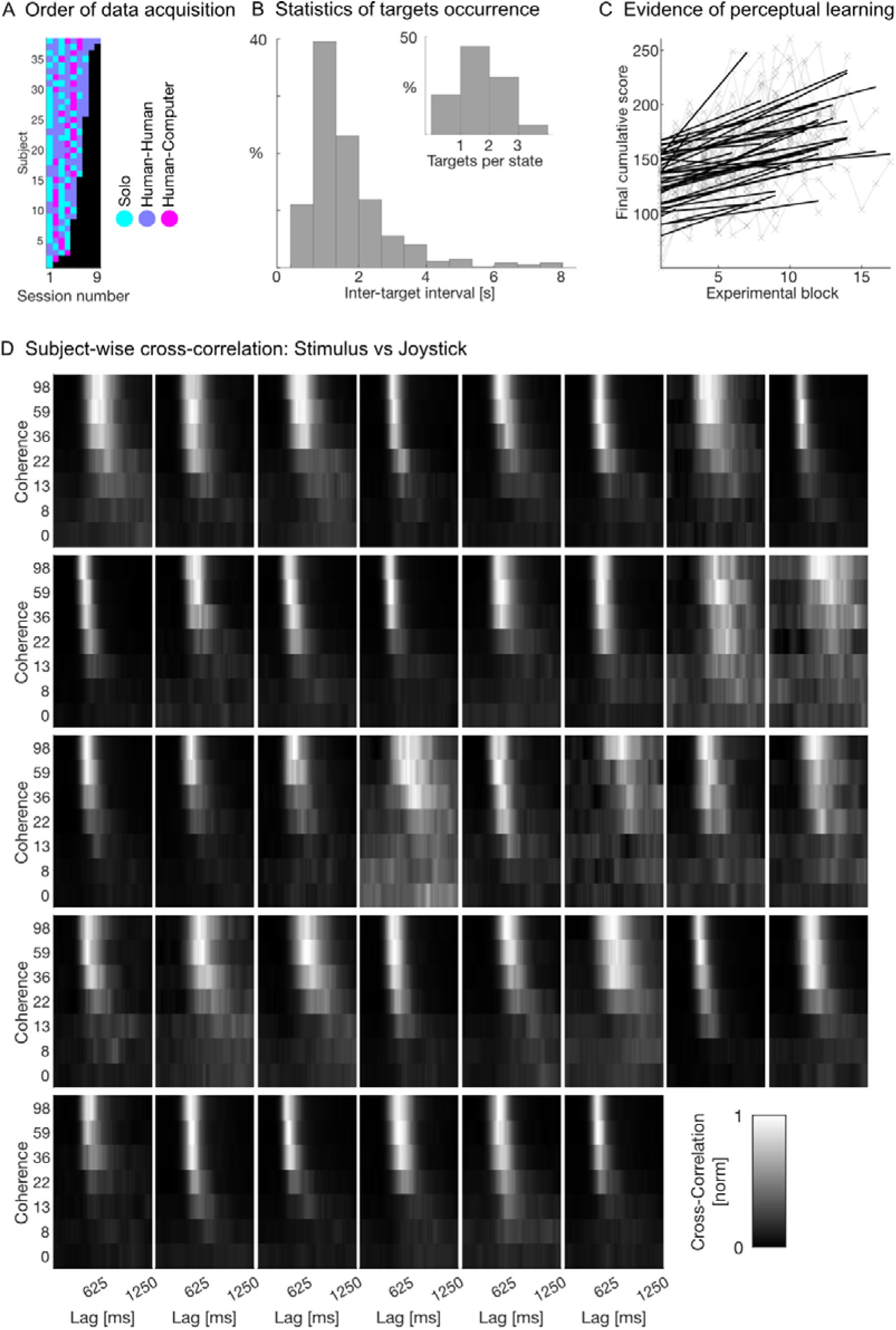
Related to Figure 1 and Figure 2. Additional information regarding experiments and joystick responses of individual participants. **(A)** Number and identity of experimental sessions for each subject. A session comprised two experimental blocks that were recorded in different setups. The order of the session type (solo or dyadic) was mixed (color-coded). All participants, except two, contributed data to each session type, specifically, solo CPR as well as both dyadic conditions. **(B)** Statistics of target occurrences during stimulus presentation. Distributions of inter-target intervals and target count per stimulus state for an example session. Targets were flashed with a 1% probability every 10 ms. Once a target was presented it remained in the screen for 50 ms followed by a minimum inter-target interval of 300 ms. **(C)** Final reward scores of participants over the course of the individuals’ data acquisition period (gray lines). Reward score increased over time. A linear regression was fitted to the cumulative scores of each experimental block for each participant (black lines). Note that each experimental session comprised two blocks, one in each setup. Independent of the session type (see panel (A) for more details), scores increased over time, likely due to perceptual learning. The final cumulative score is comprised of the hit rate, accuracy and tilt, all of which are affected by number of the experimental block, with later sessions resulting in higher hit rate as well as more accurate and tilted responses (**Supplementary Table 1** - **Supplementary Table 4**). **(D)** Normalized cross-correlation coefficients between random dot motion direction and joystick response direction illustrated for each subject. Lighter hues indicate higher cross-correlation coefficients at the respective signal lag, darker hues suggest low correlation between stimulus and joystick response at that lag. Cross-correlations broke down consistently with lower stimulus coherence.

**Supplementary Figure 2:**
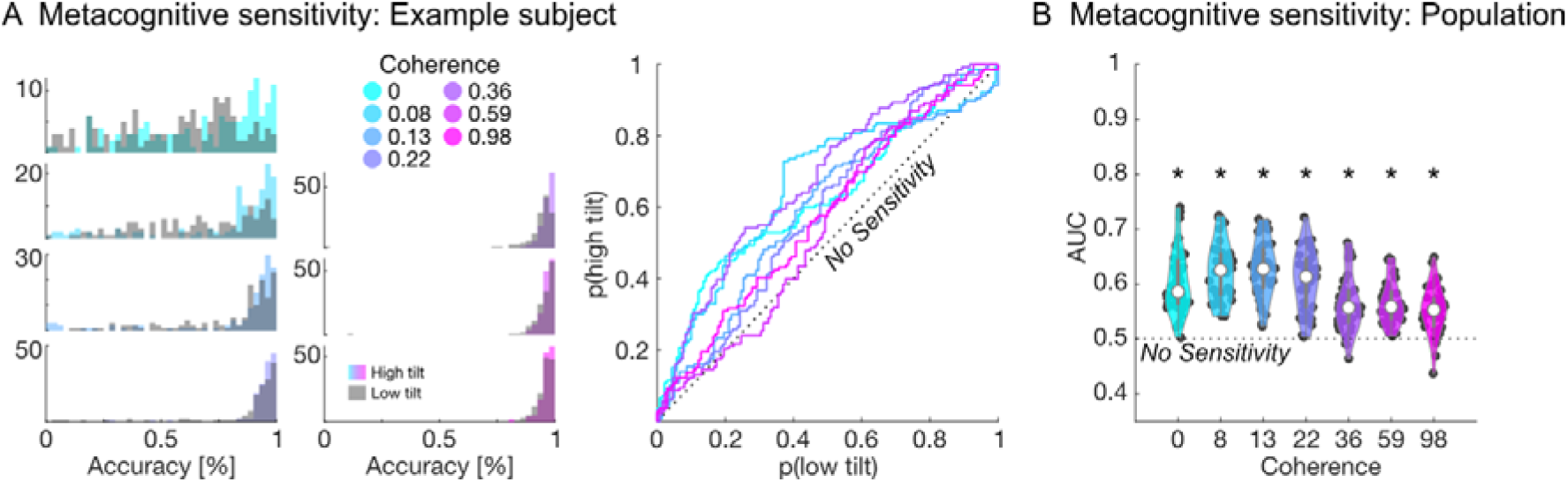
Related to Figure 2. Joystick tilt is a proxy measure of perceptual confidence. **(A)** Metacognitive sensitivity of joystick response for one example subject. Left: Distribution of joystick accuracy in stimulus states with low (gray) vs high joystick tilt (colored, median split). Accuracy and tilt were averaged for the last 30 frames (250 ms) prior to a stimulus direction change. Coherence is color-coded. Right: Corresponding receiver-operating characteristics (‘ROC’) between the two distributions for each coherence level (color-coded). A ROC curve along the diagonal would indicate similar accuracy distributions between hits and misses, suggesting no metacognitive sensitivity. **(B)** Population AUC values (black dots) are consistently above 0.5 (p<0.001, Two-sided Wilcoxon signed rank test for distribution with median 0.5), demonstrating that high tilt was more often associated with high accuracy, suggesting metacognitive-sensitive confidence readouts.

**Supplementary Figure 3:**
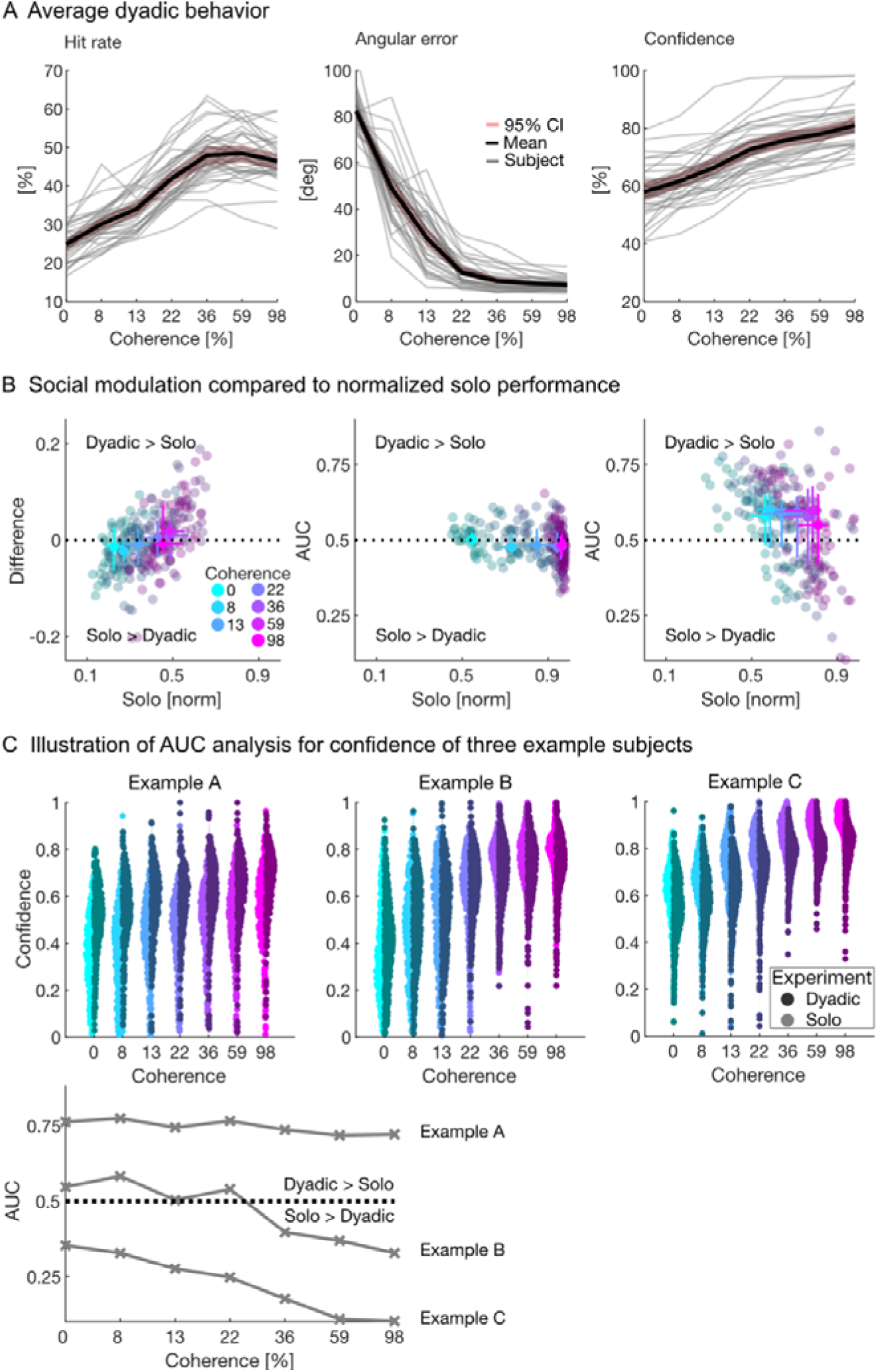
Related to Figure 3. Additional information for human-human dyadic performance. **(A)** Performance summary in (human-human) dyadic experiments. Hit rate (left), accuracy (center) and confidence (right) illustrated for each subject contributing human-human dyadic data. Same conventions as in Figure 2A. **(B)** Comparison of social modulation with response measures of solo experiments. Hit rate differences (left, compared to solo hit rate) and social modulation of accuracy (center, compared to solo accuracy) and tilt (right, compared to solo confidence) are displayed for the entire population. Joystick data was recorded in a normalized fashion for both accuracy (180deg difference = 0; Perfect match of stimulus direction = 1) and confidence (center = 0; Max. joystick tilt = 1). Each participant contributes on data point per coherence condition (color-coded). The median score across all subjects is overlaid for each coherence condition in brighter color hues. Error bars show 99% confidence intervals of the median in solo and dyadic conditions. **(C)** Social modulation of confidence for three example participants. Data for solo (light, left) and human-human dyadic experiments (dark, right) are displayed for each stimulus coherence level (color-coded). Each dot corresponds to the time-window average for a single stimulus state. All sessions of the same experimental condition are pooled. Corresponding AUC values, used to quantify the direction and magnitude of social modulation between dyadic and solo experiments, are shown below. A value of 0.5 corresponds to perfect overlap between solo and dyadic response distributions, 1 and 0 imply perfect separation between experimental conditions.

**Supplementary Figure 4:**
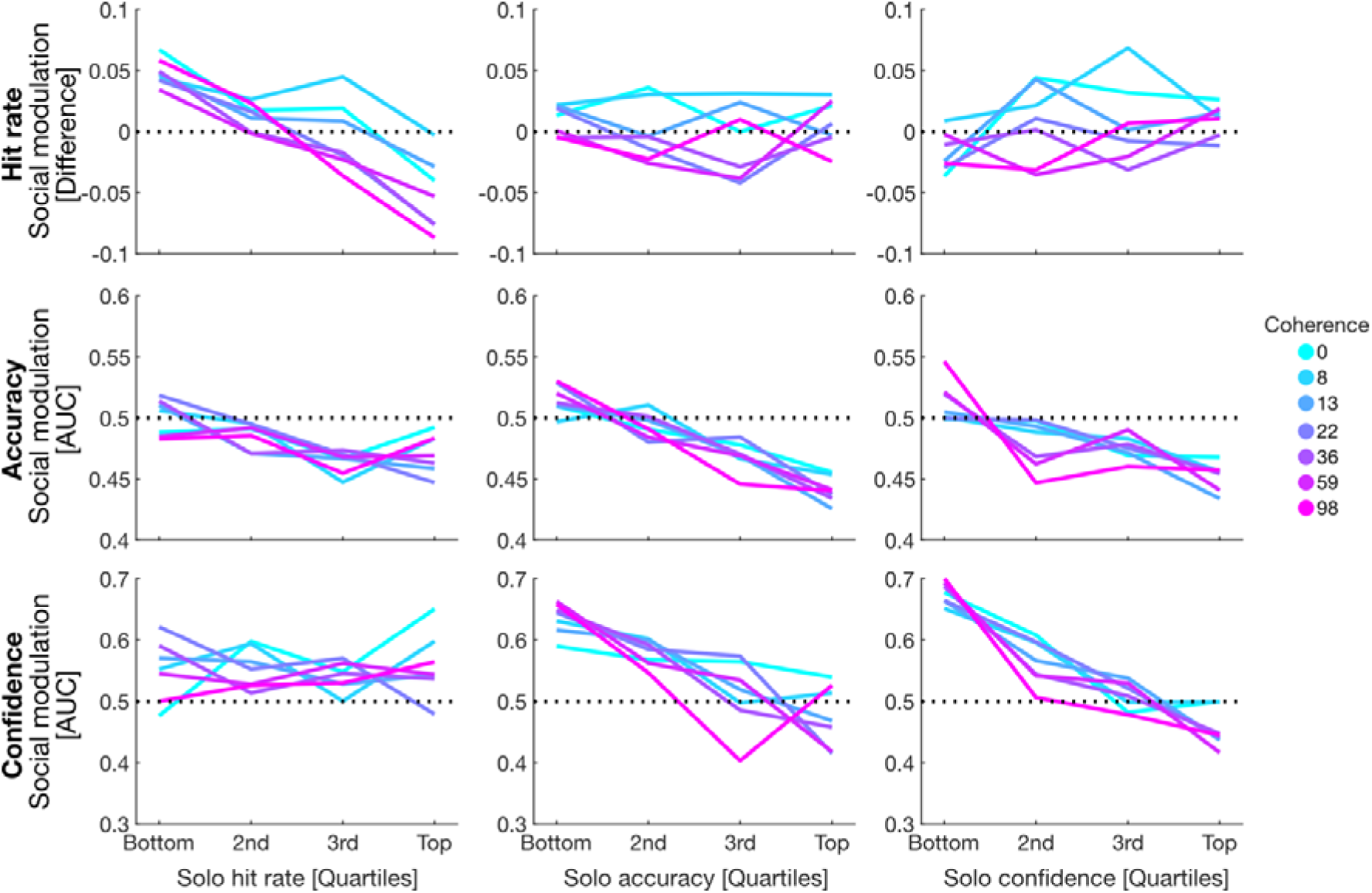
Related to Figure 3. Social modulation grouped by different metrics. Illustration of average social modulation for hit rates (first row), accuracy (second row) and confidence (third row) based on differently grouped solo performance quartiles: first column - hit rate quartiles, second column - accuracy quartiles, third column - confidence quartiles. Coherence is color-coded. Same conventions as in Figure 3C. Social modulation between dyadic and solo experiments was measured by AUC. An AUC value of 0.5 corresponds to perfect overlap between solo and dyadic response distributions. AUC > 0.5 imply better accuracy or higher confidence in dyadic experiments. AUC < 0.5 imply better accuracy or higher confidence in solo experiments.

**Supplementary Figure 5:**
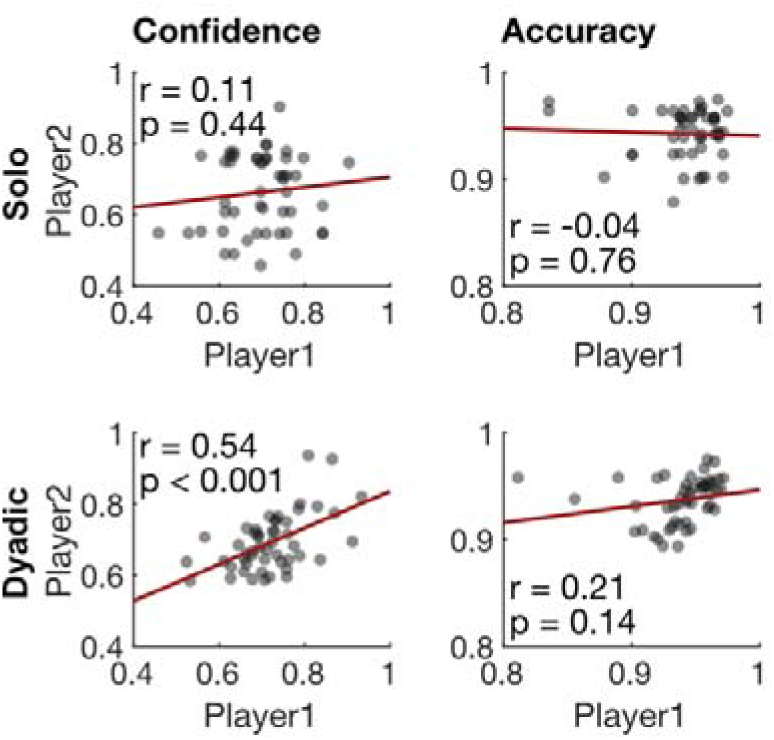
Related to Figure 4. Across-player correlation in solo and dyadic setting. Confidence (left) and accuracy (right) of dyadic partners in solo and dyadic settings. Only in the dyadic condition confidence (but not accuracy) correlate significantly between players.

**Supplementary Figure 6:**
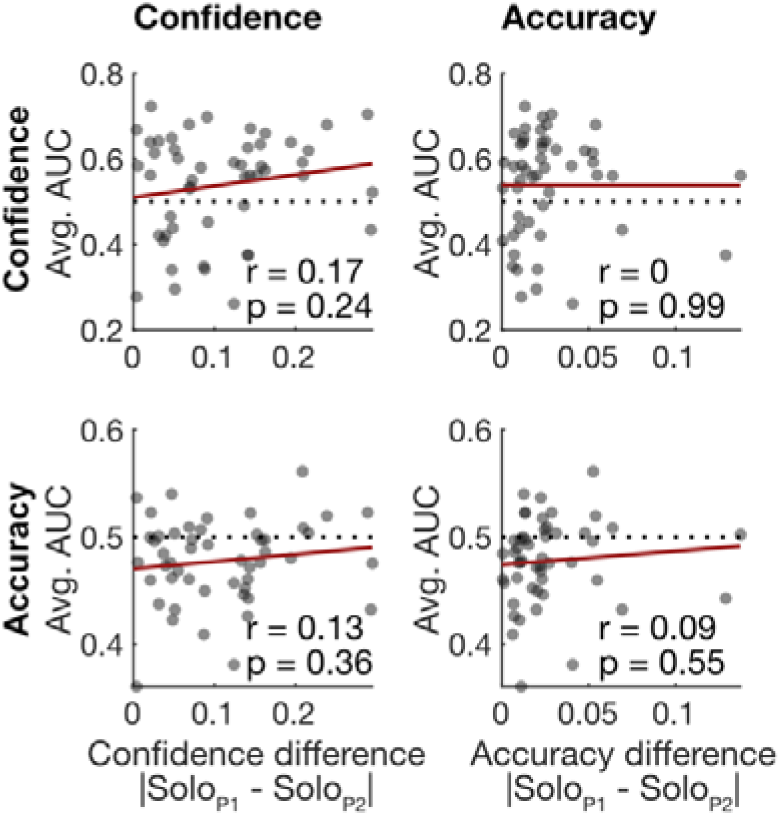
Related to Figure 4. Relationship between social modulation and solo performance difference. Average (across dyad members) social modulation for confidence (top) and accuracy (bottom) displayed as a function of the absolute solo differences in confidence (left) and accuracy (right) between dyadic partners. Each data point corresponds to one dyad (N=50). Average social modulation did not correlate with absolute solo difference between players.

**Supplementary Figure 7:**
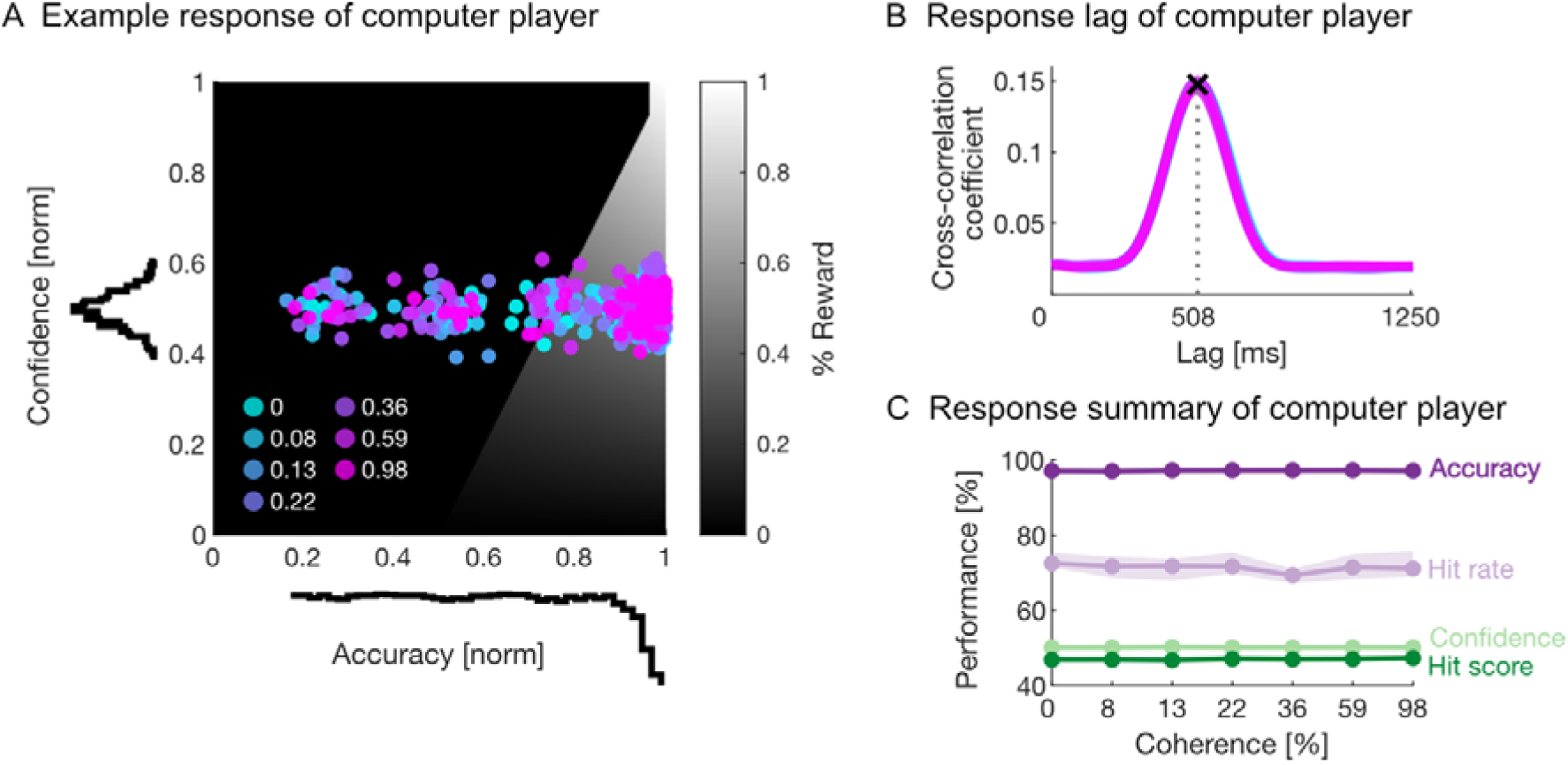
Related to Figure 5. Additional information regarding computer player performance. **(A)** Illustration of computer player behavior in an example session. Each dot corresponds to the accuracy-confidence combination during target presentation. Distribution of computer behavior is summarized with histograms. Same conventions as in Figure 1D. Coherence is color-coded. **(B)** Cross-correlation between stimulus direction changes and cursor responses of computer player. Similar human-like response lag was built in for all coherence conditions (color-coded). See Figure 2C for comparison. **(C)** Average computer player performance (reward score, hit rate, accuracy, and confidence) as a function of stimulus coherence. Shaded background corresponds to the 99% confidence intervals of the median. See Figure 2A for comparison to human behavior.

**Supplementary Figure 8:**
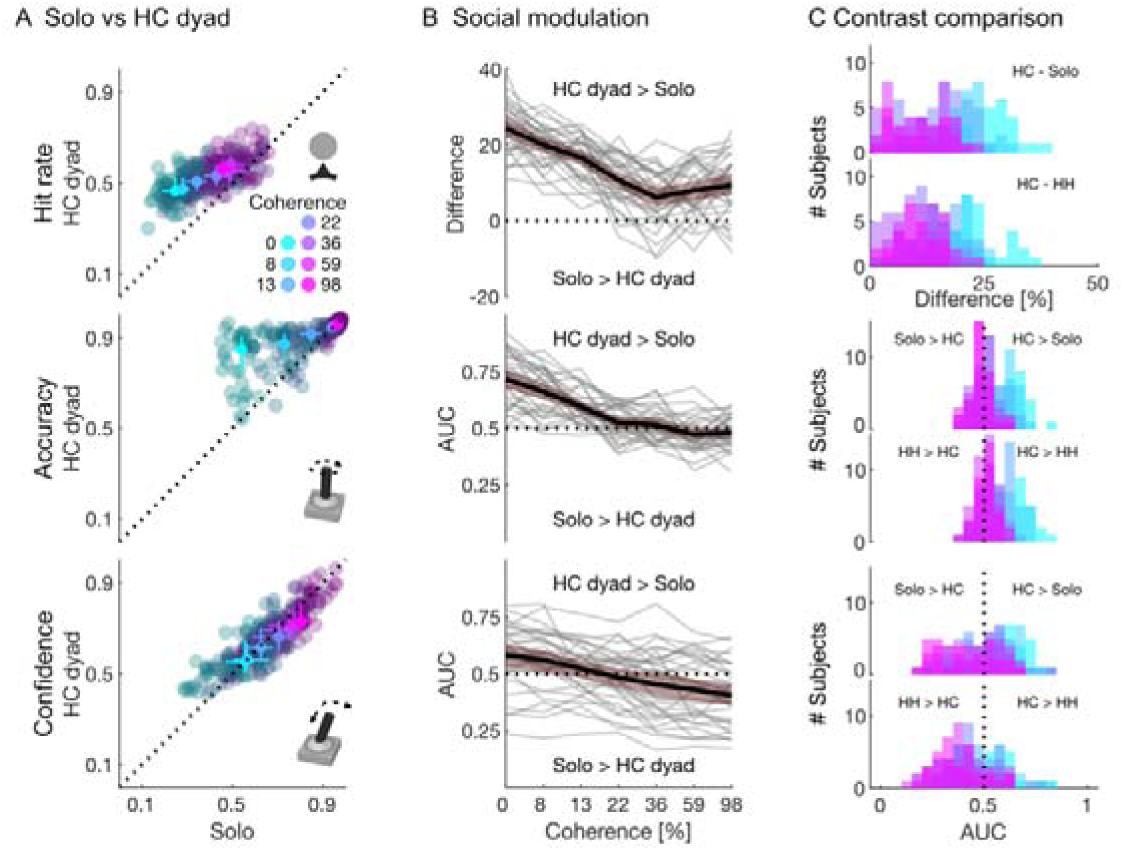
Related to Figure 5. Social modulation in human-computer (HC) dyads vs solo. **(A)** Comparison of average hit rate (top), accuracy (center) and confidence (bottom) for each stimulus coherence in solo and human-computer dyads, color-coded for coherence level. Individual data is shown in darker hues. Each subject contributes one data point per coherence condition. Medians across subjects overlaid for each coherence condition (bright color). Error bars show 99% confidence intervals of the median. Same conventions as in Figure 5. **(B)** Social modulation of humans between dyadic (human-computer, HC) and solo experiments. **(C)** Population comparison of social modulation (between solo and HH experiments, top) and HH and HC contrast (bottom).

## Supplementary Tables

**Supplementary Table 1:**
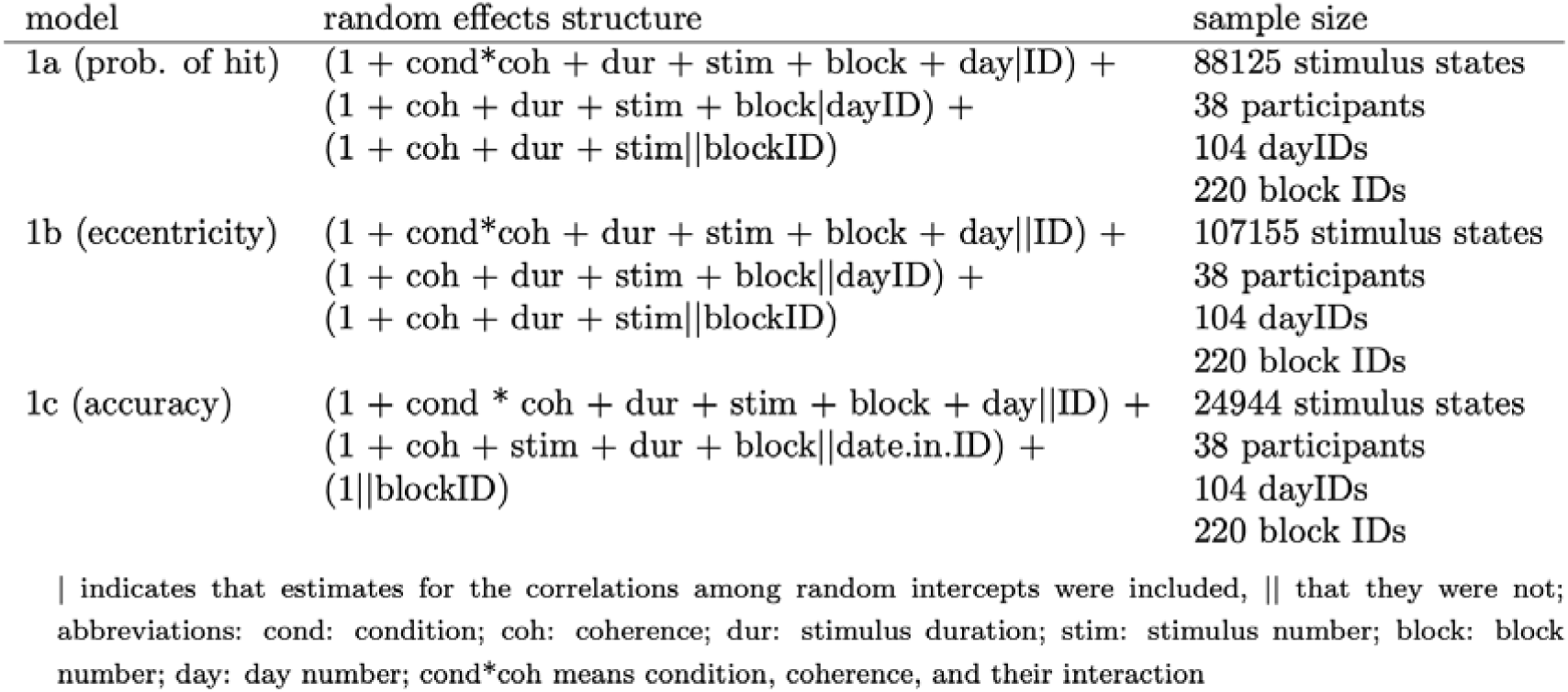
Random effects structure and sample size of each model.

**Supplementary Table 2:** Results of the full model with hit probability as response.

| term | estimate | SE | CL<sub>lower</sub> | CL<sub>upper</sub> | $\chi^2$ | df | P | min | max |
| --- | --- | --- | --- | --- | --- | --- | --- | --- | --- |
| intercept | 0.505 | 0.017 | 0.473 | 0.540 |  |  |  | 0.498 | 0.510 |
| condition | 0.238 | 0.015 | 0.208 | 0.266 |  |  |  | 0.230 | 0.244 |
| coherence | 0.135 | 0.009 | 0.118 | 0.154 |  |  |  | 0.131 | 0.139 |
| stim. duration | 0.076 | 0.005 | 0.066 | 0.085 | 91.707 | 1 | <0.001 | 0.074 | 0.077 |
| stim. nr. | 0.008 | 0.005 | -0.002 | 0.017 | 4.532 | 1 | 0.033 | 0.006 | 0.009 |
| block nr. | 0.016 | 0.006 | 0.005 | 0.028 | 6.466 | 1 | 0.011 | 0.013 | 0.020 |
| day nr. | 0.017 | 0.008 | 0.001 | 0.032 | 3.862 | 1 | 0.049 | 0.011 | 0.020 |
| condition:coherence | -0.064 | 0.011 | -0.087 | -0.041 | 26.472 | 1 | <0.001 | -0.067 | -0.061 |

indicated are estimates, together with their standard errors, confidence limits, significance tests, and the range of estimates when excluding individuals one at a time; all covariates were z-transformed, mean and standard of the original variables were 0.338 and 0.320 (coherence), 246.129 and 142.150 (stim. nr.), 1907.127 and 358.333 (stim. dur.), 1.669 and 0.754 (block nr.), and 2.982 and 1.526 (day nr.) respectively; condition was dummy coded with CPRsolo being the reference level

**Supplementary Table 3:**
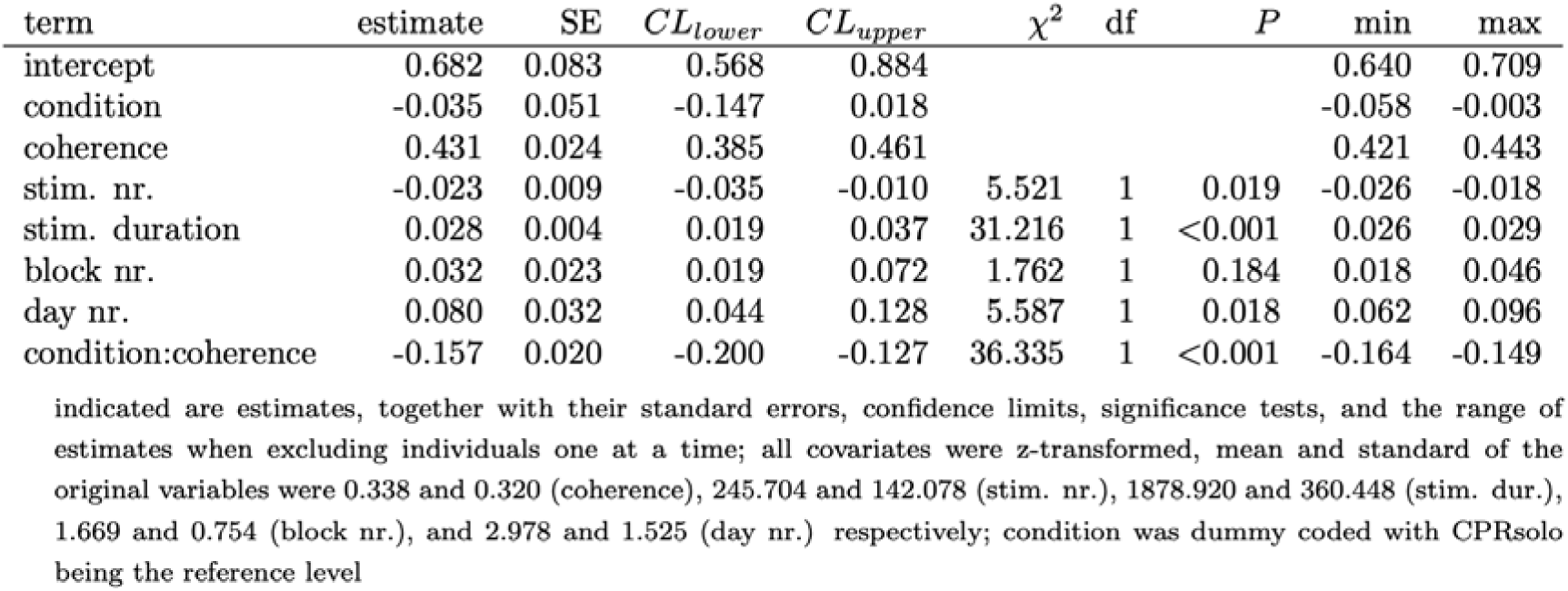
Results of the full model with confidence being the response.

| term | estimate | SE | $CL_{lower}$ | $CL_{upper}$ | $\chi^2$ | df | $P$ | min | max |
| --- | --- | --- | --- | --- | --- | --- | --- | --- | --- |
| intercept | 0.682 | 0.083 | 0.568 | 0.884 |  |  |  | 0.640 | 0.709 |
| condition | -0.035 | 0.051 | -0.147 | 0.018 |  |  |  | -0.058 | -0.003 |
| coherence | 0.431 | 0.024 | 0.385 | 0.461 |  |  |  | 0.421 | 0.443 |
| stim. nr. | -0.023 | 0.009 | -0.035 | -0.010 | 5.521 | 1 | 0.019 | -0.026 | -0.018 |
| stim. duration | 0.028 | 0.004 | 0.019 | 0.037 | 31.216 | 1 | <0.001 | 0.026 | 0.029 |
| block nr. | 0.032 | 0.023 | 0.019 | 0.072 | 1.762 | 1 | 0.184 | 0.018 | 0.046 |
| day nr. | 0.080 | 0.032 | 0.044 | 0.128 | 5.587 | 1 | 0.018 | 0.062 | 0.096 |
| condition:coherence | -0.157 | 0.020 | -0.200 | -0.127 | 36.335 | 1 | <0.001 | -0.164 | -0.149 |

indicated are estimates, together with their standard errors, confidence limits, significance tests, and the range of estimates when excluding individuals one at a time; all covariates were z-transformed, mean and standard of the original variables were 0.338 and 0.320 (coherence), 245.704 and 142.078 (stim. nr.), 1878.920 and 360.448 (stim. dur.), 1.669 and 0.754 (block nr.), and 2.978 and 1.525 (day nr.) respectively; condition was dummy coded with CPRsolo being the reference level

**Supplementary Table 4:** Results of the full model with accuracy being the response.

| term | estimate | SE | $CL_{lower}$ | $CL_{upper}$ | $\chi^2$ | df | $P$ | min | max |
| --- | --- | --- | --- | --- | --- | --- | --- | --- | --- |
| intercept | 1.354 | 0.035 | 1.283 | 1.418 |  |  |  | 1.345 | 1.373 |
| condition | 0.208 | 0.028 | 0.157 | 0.262 |  |  |  | 0.200 | 0.219 |
| coherence | 0.519 | 0.013 | 0.493 | 0.541 |  |  |  | 0.515 | 0.529 |
| stim. nr. | 0.010 | 0.007 | -0.004 | 0.023 | 2.249 | 1 | 0.134 | 0.009 | 0.015 |
| stim. duration | 0.026 | 0.006 | 0.013 | 0.039 | 16.033 | 1 | <0.001 | 0.024 | 0.029 |
| block nr. | 0.040 | 0.007 | 0.026 | 0.055 | 22.652 | 1 | <0.001 | 0.038 | 0.044 |
| day nr. | 0.060 | 0.012 | 0.036 | 0.084 | 14.045 | 1 | <0.001 | 0.053 | 0.066 |
| condition:coherence | -0.251 | 0.024 | -0.294 | -0.203 | 51.005 | 1 | <0.001 | -0.260 | -0.244 |

indicated are estimates, together with their standard errors, confidence limits, significance tests, and the range of estimates when excluding individuals one at a time; all covariates were z-transformed, mean and standard of the original variables were 0.339 and 0.321 (coherence), 245.815 and 142.924 (stim. nr.), 1979.819 and 334.555 (stim. dur.), 1.671 and 0.753 (block nr.), and 3.007 and 1.534 (day nr.) respectively; condition was dummy coded with CPRsolo being the reference level

***Supplementary Video 1:* Schematic animation of the continuous perceptual report task.** Animation is based a solo experiment of an example subject. White random dot pattern moves continuously on black background with frequently changing motion direction and coherence. The white vector indicates the nominal motion direction of the stimulus. It is scaled by motion coherence for illustration purposes. The coherence is indicated in the lower right corner. The red vector shows the behavioral response of the subject: joystick direction (polar angle) and joystick tilt (length). Both joystick response values as well as the resulting score are stated in the lower left corner. The red shading demonstrates the angular width of the response arc.

## References

Baayen RH. 2008. Analyzing Linguistic Data: A Practical Introduction to Statistics using R, 1st ed. Cambridge University Press. doi:10.1017/CBO9780511801686

Babiloni F, Astolfi L. 2014. Social neuroscience and hyperscanning techniques: Past, present and future. Neuroscience & Biobehavioral Reviews 44:76–93. doi:10.1016/j.neubiorev.2012.07.006

Bahrami B, Olsen K, Bang D, Roepstorff A, Rees G, Frith C. 2012a. What failure in collective decision-making tells us about metacognition. Philosophical Transactions of the Royal Society B: Biological Sciences 367:1350–1365. doi:10.1098/rstb.2011.0420

Bahrami B, Olsen K, Bang D, Roepstorff A, Rees G, Frith C. 2012b. Together, slowly but surely: The role of social interaction and feedback on the build-up of benefit in collective decision-making. Journal of Experimental Psychology: Human Perception and Performance 38:3–8. doi:10.1037/a0025708

Bahrami B, Olsen K, Latham PE, Roepstorff A, Rees G, Frith CD. 2010. Optimally Interacting Minds. Science 329:1081–1085. doi:10.1126/science.1185718

Balsdon T, Mamassian P, Wyart V. 2021. Separable neural signatures of confidence during perceptual decisions. eLife 10:e68491. doi:10.7554/eLife.68491

Balsdon T, Wyart V, Mamassian P. 2020. Confidence controls perceptual evidence accumulation. Nat Commun 11:1753. doi:10.1038/s41467-020-15561-w

Bang D, Aitchison L, Moran R, Herce Castanon S, Rafiee B, Mahmoodi A, Lau JYF, Latham PE, Bahrami B, Summerfield C. 2017. Confidence matching in group decision-making. Nat Hum Behav 1:0117. doi:10.1038/s41562-017-0117

Bang D, Frith CD. 2017. Making better decisions in groups. R Soc open sci 4:170193. doi:10.1098/rsos.170193

Barr DJ, Levy R, Scheepers C, Tily HJ. 2013. Random effects structure for confirmatory hypothesis testing: Keep it maximal. Journal of Memory and Language 68:255–278. doi:10.1016/j.jml.2012.11.001

Bates D, Mächler M, Bolker B, Walker S. 2015. Fitting Linear Mixed-Effects Models Using lme4. J Stat Soft 67. doi:10.18637/jss.v067.i01

Baumgart KG, Byvshev P, Sliby A, Strube A, König P, Wahn B. 2020. Neurophysiological correlates of collective perceptual decision-making. Eur J of Neuroscience 51:1676–1696. doi:10.1111/ejn.14545

Bayliss AP, Frischen A, Fenske MJ, Tipper SP. 2007. Affective evaluations of objects are influenced by observed gaze direction and emotional expression q.

Boldt A, Yeung N. 2015. Shared Neural Markers of Decision Confidence and Error Detection. J Neurosci 35:3478–3484. doi:10.1523/JNEUROSCI.0797-14.2015

Bolker BM. 2008. Ecological models and data in R. Princeton, N.J: Princeton University Press.

Bonnen K, Burge J, Yates J, Pillow J, Cormack LK. 2015. Continuous psychophysics: Target-tracking to measure visual sensitivity. Journal of Vision 15:14. doi:10.1167/15.3.14

Bonnen K, Huk AC, Cormack LK. 2017. Dynamic mechanisms of visually guided 3D motion tracking. Journal of Neurophysiology 118:1515–1531. doi:10.1152/jn.00831.2016

Brooks M E, Kristensen K, Benthem K J, van, Magnusson A, Berg C W, Nielsen A, Skaug H J, Mächler M, Bolker B M. 2017. glmmTMB Balances Speed and Flexibility Among Packages for Zero-inflated Generalized Linear Mixed Modeling. The R Journal 9:378. doi:10.32614/RJ-2017-066

Czeszumski A, Eustergerling S, Lang A, Menrath D, Gerstenberger M, Schuberth S, Schreiber F, Rendon ZZ, König P. 2020. Hyperscanning: A Valid Method to Study Neural Inter-brain Underpinnings of Social Interaction. Front Hum Neurosci 14:39. doi:10.3389/fnhum.2020.00039

De Martino B, Bobadilla-Suarez S, Nouguchi T, Sharot T, Love BC. 2017. Social Information Is Integrated into Value and Confidence Judgments According to Its Reliability. J Neurosci 37:6066–6074. doi:10.1523/JNEUROSCI.3880-16.2017

Dobson A. 2001. An Introduction to Generalized Linear Models, Second Edition, Chapman & Hall/CRC Texts in Statistical Science. Chapman and Hall/CRC. doi:10.1201/9781420057683

Dosso JA, Roberts KH, DiGiacomo A, Kingstone A. 2018. The Influence of Co-action on a Simple Attention Task: A Shift Back to the Status Quo. Front Psychol 9:874. doi:10.3389/fpsyg.2018.00874

Esmaily J, Zabbah S, Ebrahimpour R, Bahrami B. 2023. Interpersonal alignment of neural evidence accumulation to social exchange of confidence. eLife 12:e83722. doi:10.7554/eLife.83722

Fan Y, Gold JI, Ding L. 2018. Ongoing, rational calibration of reward-driven perceptual biases. eLife 7:1–26. doi:10.7554/eLife.36018

Farrell S. 2011. Social influence benefits the wisdom of individuals in the crowd. Proc Natl Acad Sci USA 108. doi:10.1073/pnas.1109947108

Fetsch CR, Kiani R, Newsome WT, Shadlen MN. 2014. Effects of Cortical Microstimulation on Confidence in a Perceptual Decision. Neuron 83:797–804. doi:10.1016/j.neuron.2014.07.011

Fetsch CR, Odean NN, Jeurissen D, El-Shamayleh Y, Horwitz GD, Shadlen MN. 2018. Focal optogenetic suppression in macaque area MT biases direction discrimination and decision confidence, but only transiently. eLife 7:1–23. doi:10.7554/eLife.36523

Fleming SM, Lau HC. 2014. How to measure metacognition. Frontiers in Human Neuroscience 8:1–9. doi:10.3389/fnhum.2014.00443

Fleming SM, Weil RS, Nagy Z, Dolan RJ, Rees G. 2010. Relating Introspective Accuracy to Individual Differences in Brain Structure. Science 329:1541–1543. doi:10.1126/science.1191883

Forstmeier W, Schielzeth H. 2011. Cryptic multiple hypotheses testing in linear models: overestimated effect sizes and the winner’s curse. Behav Ecol Sociobiol 65:47–55. doi:10.1007/s00265-010-1038-5

Frith CD, Singer T. 2008. The role of social cognition in decision making. Phil Trans R Soc B 363:3875–3886. doi:10.1098/rstb.2008.0156

Gail A, Brinksmeyer, H. J., Eckhorn, R. 2004. Perception-related Modulations of Local Field Potential Power and Coherence in Primary Visual Cortex of Awake Monkey during Binocular Rivalry. Cerebral Cortex 14:300–313. doi:10.1093/cercor/bhg129

Germar M, Albrecht T, Voss A, Mojzisch A. 2016. Social conformity is due to biased stimulus processing: electrophysiological and diffusion analyses. Social Cognitive and Affective Neuroscience 11:1449–1459. doi:10.1093/scan/nsw050

Gold JI, Stocker AA. 2017. Visual Decision-Making in an Uncertain and Dynamic World. Annu Rev Vis Sci 3:227–250. doi:10.1146/annurev-vision-111815-114511

Hanks TD, Summerfield C. 2017. Perceptual Decision Making in Rodents, Monkeys, and Humans. Neuron 93:15–31. doi:10.1016/j.neuron.2016.12.003

Huk A, Bonnen K, He BJ. 2018. Beyond Trial-Based Paradigms: Continuous Behavior, Ongoing Neural Activity, and Natural Stimuli. J Neurosci 38:7551–7558. doi:10.1523/JNEUROSCI.1920-17.2018

Isbaner S, Mendoza RB, Bothe R, Fischer A, Gail A, Gläscher J, Lueschen HS, Moeller S, Eiteljoerge SFV, Penke L, Priesemann V, Ruß J, Schacht A, Schneider F, Shahidi N, Treue S, Wibral M, Ziereis A, Fischer J, Kagan I, Mani N. 2025. Dyadic Interaction Platform: A novel tool to study transparent social interactions. doi:10.31234/osf.io/2rckn_v1

Kepecs A, Mainen ZF. 2012. A computational framework for the study of confidence in humans and animals. Phil Trans R Soc B 367:1322–1337. doi:10.1098/rstb.2012.0037

Khalvati K, Kiani R, Rao RPN. 2021. Bayesian inference with incomplete knowledge explains perceptual confidence and its deviations from accuracy. Nat Commun 12:5704. doi:10.1038/s41467-021-25419-4

Kiani R, Shadlen MN. 2009. Representation of Confidence Associated with a Decision by Neurons in the Parietal Cortex. Science 324:759–764. doi:10.1126/science.1169405

Komura Y, Nikkuni A, Hirashima N, Uetake T, Miyamoto A. 2013. Responses of pulvinar neurons reflect a subject’s confidence in visual categorization. nature NEUROSCIENCE.

Kruger J, Dunning D. 1999. Unskilled and unaware of it: How difficulties in recognizing one’s own incompetence lead to inflated self-assessments. Journal of Personality and Social Psychology 77:1121–1134. doi:10.1037/0022-3514.77.6.1121

Lewen D, Ivanov V, Dehning J, Ruß J, Fischer A, Penke L, Schacht A, Gail A, Priesemann V, Kagan I. 2025. Continuous dynamics of cooperation and competition in social foraging. doi:10.1101/2025.05.28.655569

Lorenz J, Rauhut H, Schweitzer F, Helbing D. 2011. How social influence can undermine the wisdom of crowd effect. Proc Natl Acad Sci USA 108:9020–9025. doi:10.1073/pnas.1008636108

Madipakkam AR, Bellucci G, Rothkirch M, Park SQ. 2019. The influence of gaze direction on food preferences. Sci Rep 9:5604. doi:10.1038/s41598-019-41815-9

Mahmoodi A, Bang D, Olsen K, Zhao YA, Shi Z, Broberg K, Safavi S, Han S, Nili Ahmadabadi M, Frith CD, Roepstorff A, Rees G, Bahrami B. 2015. Equality bias impairs collective decision-making across cultures. Proc Natl Acad Sci USA 112:3835–3840. doi:10.1073/pnas.1421692112

Maniscalco B, Lau H. 2014. Signal Detection Theory Analysis of Type 1 and Type 2 Data: Meta-d1, Response-Specific Meta-d1, and the Unequal Variance SDT Model In: Fleming SM, Frith CD, editors. The Cognitive Neuroscience of Metacognition. Berlin, Heidelberg: Springer Berlin Heidelberg. pp. 25–66. doi:10.1007/978-3-642-45190-4_3

Maniscalco B, Lau H. 2012. A signal detection theoretic approach for estimating metacognitive sensitivity from confidence ratings. Consciousness and Cognition 21:422–430. doi:10.1016/j.concog.2011.09.021

Matuschek H, Kliegl R, Vasishth S, Baayen H, Bates D. 2017. Balancing Type I error and power in linear mixed models. Journal of Memory and Language 94:305–315. doi:10.1016/j.jml.2017.01.001

McCullagh P, Nelder JA. 1989. Generalized Linear Models. Boston, MA: Springer US. doi:10.1007/978-1-4899-3242-6

Mitsuda T, Masaki S. 2018. Subliminal gaze cues increase preference levels for items in the gaze direction. Cognition and Emotion 32:1146–1151. doi:10.1080/02699931.2017.1371002

Moeller S, Unakafov AM, Fischer J, Gail A, Treue S, Kagan I. 2023. Human and macaque pairs employ different coordination strategies in a transparent decision game. eLife 12:e81641. doi:10.7554/eLife.81641

Moreira CM, Rollwage M, Kaduk K, Wilke M, Kagan I. 2018. Post-decision wagering after perceptual judgments reveals bi-directional certainty readouts. Cognition 176:40–52. doi:10.1016/j.cognition.2018.02.026

Navajas J, Bahrami B, Latham PE. 2016. Post-decisional accounts of biases in confidence. Current Opinion in Behavioral Sciences 11:55–60. doi:10.1016/j.cobeha.2016.05.005

Noel J-P, Balzani E, Avila E, Lakshminarasimhan KJ, Bruni S, Alefantis P, Savin C, Angelaki DE. 2022. Coding of latent variables in sensory, parietal, and frontal cortices during closed-loop virtual navigation. eLife 11:e80280. doi:10.7554/eLife.80280

Noel J-P, Bill J, Ding H, Vastola J, DeAngelis GC, Angelaki DE, Drugowitsch J. 2023. Causal inference during closed-loop navigation: parsing of self- and object-motion. Phil Trans R Soc B 378:20220344. doi:10.1098/rstb.2022.0344

Park SA, Goïame S, O’Connor DA, Dreher J-C. 2017. Integration of individual and social information for decision-making in groups of different sizes. PLoS Biol 15:e2001958. doi:10.1371/journal.pbio.2001958

Persaud N, McLeod P, Cowey A. 2007. Post-decision wagering objectively measures awareness. Nat Neurosci 10:257–261. doi:10.1038/nn1840

Pescetelli N, Rees G, Bahrami B. 2016. The perceptual and social components of metacognition. Journal of Experimental Psychology: General 145:949–965. doi:10.1037/xge0000180

Pescetelli N, Yeung N. 2022. Benefits of spontaneous confidence alignment between dyad members. Collective Intelligence 1:263391372211269. doi:10.1177/26339137221126915

Pescetelli N, Yeung N. 2020. The effects of recursive communication dynamics on belief updating. Proc R Soc B 287:20200025. doi:10.1098/rspb.2020.0025

Rauhut H, Lorenz J, Schweitzer F, Helbing D. 2011. Reply to Farrell: Improved individual estimation success can imply collective tunnel vision. Proc Natl Acad Sci USA 108. doi:10.1073/pnas.1111007108

Salzman CD, Britten KH, Newsome WT. 1990. Cortical microstimulation influences perceptual judgements of motion direction. Nature 346:174–177. doi:10.1038/346174a0

Schielzeth H, Forstmeier W. 2009. Conclusions beyond support: overconfident estimates in mixed models. Behavioral Ecology 20:416–420. doi:10.1093/beheco/arn145

Smith J. 1997. The uncertain response in humans and animals. Cognition 62:75–97. doi:10.1016/S0010-0277(96)00726-3

Smithson M, Verkuilen J. 2006. A better lemon squeezer? Maximum-likelihood regression with beta-distributed dependent variables. Psychological Methods 11:54–71. doi:10.1037/1082-989X.11.1.54

Straub D, Rothkopf CA. 2022. Putting perception into action with inverse optimal control for continuous psychophysics. eLife 11. doi:10.7554/eLife.76635

Szul MJ, Bompas A, Sumner P, Zhang J. 2020. The validity and consistency of continuous joystick response in perceptual decision-making. Behav Res 52:681–693. doi:10.3758/s13428-019-01269-3

Takagaki K, Krug K. 2020. The effects of reward and social context on visual processing for perceptual decision-making. Current Opinion in Physiology 16:109–117. doi:10.1016/j.cophys.2020.08.006

Terenzi D, Liu L, Bellucci G, Park SQ. 2021. Determinants and modulators of human social decisions. Neuroscience & Biobehavioral Reviews 128:383–393. doi:10.1016/j.neubiorev.2021.06.041

Toelch U, Dolan RJ. 2015. Informational and Normative Influences in Conformity from a Neurocomputational Perspective. Trends in Cognitive Sciences 19:579–589. doi:10.1016/j.tics.2015.07.007

Toelch U, Pooresmaeili A, Dolan RJ. 2018. Neural substrates of norm compliance in perceptual decisions. Sci Rep 8:3315. doi:10.1038/s41598-018-21583-8

Van Den Bos R, Jolles JW, Homberg JR. 2013. Social modulation of decision-making: a cross-species review. Front Hum Neurosci 7. doi:10.3389/fnhum.2013.00301

Yeung N, Summerfield C. 2012. Metacognition in human decision-making: confidence and error monitoring. Phil Trans R Soc B 367:1310–1321. doi:10.1098/rstb.2011.0416

